# Data-driven predictive design of engineered living hydrogels

**DOI:** 10.64898/2026.09.17.752317

**Authors:** Geisler Muñoz-Guamuro, Payman Goodarzi, Sadaf Reihani, Roland Bennewitz, Viktor Zaverkin, Wilfried Weber

## Abstract

Engineered living materials (ELMs) offer a promising route to biologically manufactured materials for healthcare, construction and manufacturing. However, their rational design is limited by the lack of quantitative relationships linking design parameters to material properties. Here, we show that a tabular foundation model (TabPFN), informed by a small library of living hydrogels, accurately predicts macroscopic material properties from genetic and process parameters. Models were evaluated for predicting storage modulus (*G′*), fibre content, thickness, and permeability of *Escherichia coli*-produced living hydrogels containing CsgA-based fibres fused to genetically encoded PEG-like biopolymers. On an independent validation set, TabPFN achieved the strongest prediction for *G′* (R^2^ = 85.1%), reducing RMSE by 48.0% compared to linear regression. Property-guided design further enabled identification of parameters for achieving living hydrogels with desired properties. These results establish a broadly applicable framework for predictive design of ELMs, reducing experimental screening and accelerating the discovery of materials with targeted properties.

## 1. Introduction

Advanced materials are increasingly expected to be sustainably manufactured, environmentally responsive and capable of self-maintenance. Living organisms naturally achieve these functions by assembling complex structures from simple feedstocks, adapting to changing conditions and continuously regenerating their components. Inspired by these capabilities, engineered living materials (ELMs) harness genetically programmable cells as active components of material systems, enabling materials that can grow, self-assemble, sense environmental signals, and produce functional biomolecules. Such systems have attracted considerable interest for applications ranging from biosensing and environmental remediation to healthcare and sustainable manufacturing^1–8^. Among ELM platforms, microbial systems are particularly attractive because they can be engineered with high precision and cultivated at scale from renewable resources. By programming cellular functions and extracellular matrix production, engineered microorganisms can generate materials with tunable properties through the manipulation of genetic and cultivation parameters^4,9,7,10,11^.

Despite this potential, the rational development of ELMs remains limited by a lack of quantitative relationships between genetic and process design parameters and resulting material properties. Existing studies have established that genetic modifications and cultivation conditions can alter material composition, mechanics and function^9–11,7,12,4,13–17^. However, identifying the combination of genetic and process design parameters leading to a specific property mainly remains an empirical process. This limitation is particularly important for mechanical properties, which determine the suitability of living materials for applications ranging from soft tissue interfaces and mechanobiological systems to coatings, packaging and structural materials. Without predictive genotype and process-to-property relationships, the design space must be explored through iterative construction, cultivation and characterization, restricting both the range of materials that can be investigated and the pace at which application-specific properties can be achieved.

Establishing such relationships is challenging because the macroscopic properties of living materials emerge from interacting processes across multiple length and time scales. Genetic changes can affect gene expression, protein folding, secretion and extracellular assembly, while process conditions simultaneously influence cellular growth, matrix production, hydration and supramolecular organization. These processes can interact nonlinearly, such that neither sequence nor process parameters alone provide a direct predictor of bulk material behaviour. Although mechanistic models can describe selected aspects of gene expression, metabolism or population dynamics^18^, constructing a mechanistic model that relates genetic and process inputs to material-scale mechanics would require quantitative knowledge of numerous coupled processes that is currently unavailable.

Data-driven models offer an alternative by learning input-to-property relationships directly from experimental observations without requiring a complete description of the underlying mechanisms. Linear regression can identify simple trends, whereas nonlinear methods such as Gaussian process regression (GPR) can represent more complex relationships and provide predictive uncertainty estimates^19,20^. Their application to ELMs is nevertheless constrained by the limited amount of available data. Each observation may require genetic construction, cultivation over several days and physical characterization, making exhaustive sampling of a multidimensional design space impractical. Small experimental datasets also increase the risk of overfitting and often preclude the use of conventional deep-learning models.

Recent developments in tabular foundation models provide a potential strategy for learning under these data-limited conditions. The Tabular Prior-data Fitted Network, TabPFN, is a transformer-based model pretrained on millions of synthetic datasets generated from diverse data-generating processes^21^. When applied to a new prediction task, TabPFN uses the available training observations as context and draws on statistical patterns acquired during pretraining, reducing the need for extensive task-specific training and hyperparameter optimization. TabPFN has demonstrated strong predictive performance on small and medium-sized tabular datasets and may therefore be particularly suitable for experimentally intensive materials systems^22–24^.

Living materials grown from engineered bacterial biofilms have gained momentum enabling applications comprising biosensing, biocatalysis, therapeutic delivery, environmental remediation, and sustainable packaging materials ^4,7–9,13^. Biofilm-forming cells naturally secrete extracellular polymers that assemble into cohesive and hydrated matrices. In *Escherichia coli*, a major proteinaceous component of this matrix consists of curli nanofibres. Curli fibres are assembled predominantly from CsgA subunits that are secreted through the curli biogenesis machinery, nucleated by CsgB at the cell surface and incorporated into extracellular amyloid fibrils^25^. Because CsgA is genetically encoded and tolerates fusion to heterologous peptide and protein domains, the molecular composition of the extracellular matrix can be genetically programmed. The Biofilm-Integrated Nanofiber Display platform (BIND) established that CsgA fusions retain extracellular self-assembly while introducing functions such as adhesion, nanoparticle templating, or protein immobilization^11^. Subsequent studies have used engineered curli matrices for three-dimensional printing, therapeutic protein presentation and the production of mechanically tunable and biodegradable materials^7,12,13^.

Curli-based systems also illustrate how genetic modifications can regulate bulk material mechanics. For example, complementary SpyTag and SpyCatcher domains fused to CsgA introduce covalent inter-fibre crosslinks and approximately double the Young’s modulus of the resulting material^7^. Genetic alterations to matrix-forming proteins have similarly changed the storage modulus of autonomously assembled bacterial materials by more than an order of magnitude^9^. These examples demonstrate that material mechanics are genetically programmable. They do not, however, provide a quantitative framework for predicting mechanics from genetic and process design parameters or for identifying the design parameters required to obtain a target mechanical property.

In this study, we investigate whether data-driven models can predict the storage modulus (*G′*) of curli-based engineered living hydrogels from one genetic and one process design parameter. As the genetic variable, we used different numbers of repeats of a proline-, alanine- and serine-rich PAS sequence fused to the C terminus of CsgA (CsgA-PAS_n_). PAS polypeptides are genetically encoded, hydrophilic and conformationally disordered polymers with biophysical properties resembling those of polyethylene glycol (PEG)^26,27^, a common hydrogel matrix. Their genetically adjustable length provides a means to tune the molecular composition and organization of the biopolymer network. As the process variable, we varied the concentration of isopropyl β-D-1-thiogalactopyranoside (IPTG) to tune expression levels of the CsgA-PAS_n_ constructs. The resulting materials displayed gel-like rheological behaviour, with *G′* depending on both PAS repeat number and IPTG-mediated induction level.

We compared linear regression, GPR and TabPFN for predicting *G′* from the number of PAS repeats and IPTG concentration. Model performance was evaluated using condition-wise and induction-level-wise cross-validation and was subsequently tested using independently produced hydrogels containing previously unseen number of PAS repeats or unseen IPTG concentration. TabPFN showed the strongest overall generalization to the independent experimental data and provided predictive intervals that encompassed all measured validation values. We then applied the trained model in a grid-based inverse-design procedure to identify combinations of genetic and process design parameters predicted to yield target storage moduli. Finally, we examined the applicability of the same framework to additional structural and functional properties, including hydrogel thickness, specific curli-fibre content and macromolecular permeability. Together, this work establishes a small-data approach for linking genetic and design parameters to the properties of bacterially produced living hydrogels and for narrowing the experimental search space for materials with specified properties.

## 2. Results

### 2.1 Design and construction of an engineered living hydrogel library

To establish quantitative relationships linking genetic and process design parameters to resulting material properties, we first designed, constructed, and characterized a library of engineered living hydrogels. We next evaluated different statistical models to capture the design parameters-material properties relationships and validated the predictions by synthesizing the corresponding materials. Finally, we applied a high-performing model in an inverse design approach to identify design parameters that yield living materials with desired properties (**Fig. 1**).

**Figure 1.**
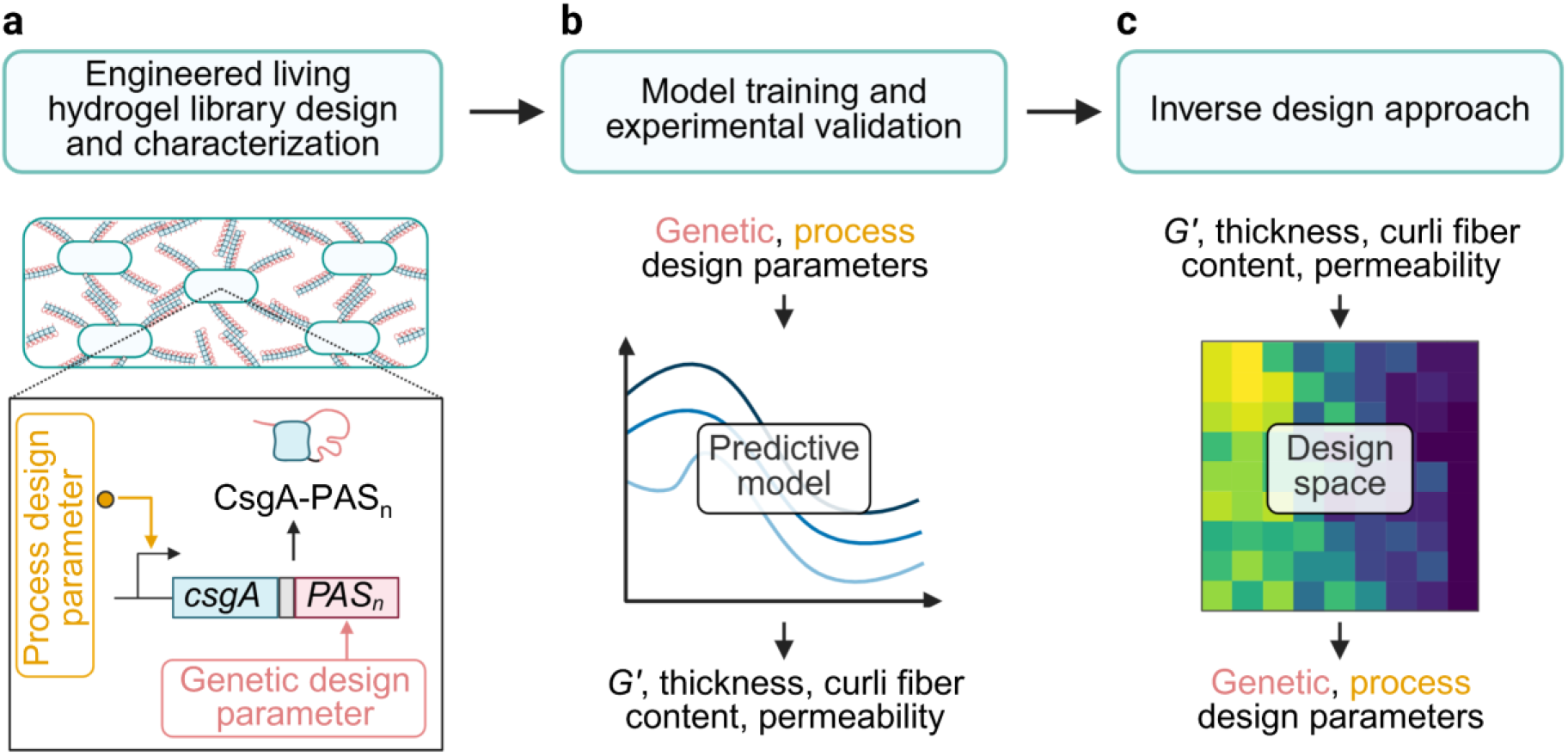
Workflow for implementing a predictive framework for the design of engineered living hydrogels. (**a**) First, we designed and built a library of curli-based living hydrogels by genetically fusing different numbers of PAS polypeptide repeats to CsgA, representing the genetic design parameter. We then collected data on the storage modulus (*G′*), thickness, curli fibre content, and permeability of the resulting living hydrogels across different concentrations of the process design parameter, isopropylthio-β-galactoside (IPTG), tuning production levels of CsgA-PAS_n_ biopolymers. (**b**) Second, we trained and cross-validated predictive models to infer material properties from these two design parameters. We further experimentally validated the models using independent, previously unseen parameter values and combinations. (**c**) Finally, using the validated models, we implemented a property-guided design approach to determine the design parameter combinations required to achieve a target material property.

We developed a library of engineered living hydrogels based on *E. coli* biofilms that could be programmed by both, genetic as well as process design parameters. As genetic design parameter, we varied the length of PAS biopolymers fused to the C-terminus of CsgA (13.3 kDa) via a three-amino acid linker (GSS). We initially used 0, 1, 5, 10 or 20 copies of the 20-amino acid core PAS motif (ASPAAPAPASPAAPAPSAPA, 1.7 kDa)^26,27^ (**Fig. 2a**). As process design parameter, we varied the concentration of isopropylthio-β-galactoside (IPTG) to tune expression strength of the CsgA-PAS_n_-encoding gene placed under the control of the P_trc_ promoter (**Fig. 2b, Supplementary Table 1, 2**). The living hydrogel variants were produced in the *E. coli* K12 MG1655-derivative PHL628 having a knockout of endogenous *csgA*^28^ and harbouring a single point mutation in the *ompR* gene which was shown to enhance curli fibre production^29^. The cells were grown on YESCA agar plates supplemented with different concentrations of the inducer IPTG (5, 15, 25 µM) for 48 h. The amount of curli nanofiber formation was estimated by staining with Congo red and subsequent image analysis for quantifying the accumulated red dye (**Fig. 2c, Supplementary Fig. 1**). We observed that the different living hydrogel variants required different IPTG concentrations for maximum curli fibre production, with 15 µM being the concentration at which the largest number of living hydrogels reached highest levels. Further, curli fibre amount remained relatively consistent across PAS repeat variants, except for the PAS_5_ variant which showed a noticeably high redness value at 5 µM IPTG, a 1.3-fold increase compared with the average redness of the other PAS variants. In general, the levels decreased with higher number of PAS repeats, which may be related to reduced secretion of the CsgA variants and assembly of curli fibre caused by the addition of the PAS polypeptides. Consistent with these observations, it has previously been reported that fusing peptides with different lengths and compositions to the C-terminus of CsgA results in distinct phenotypes on YESCA - Congo agar plates, which was attributed to different production levels of curli fibres^11^.

**Figure 2.**
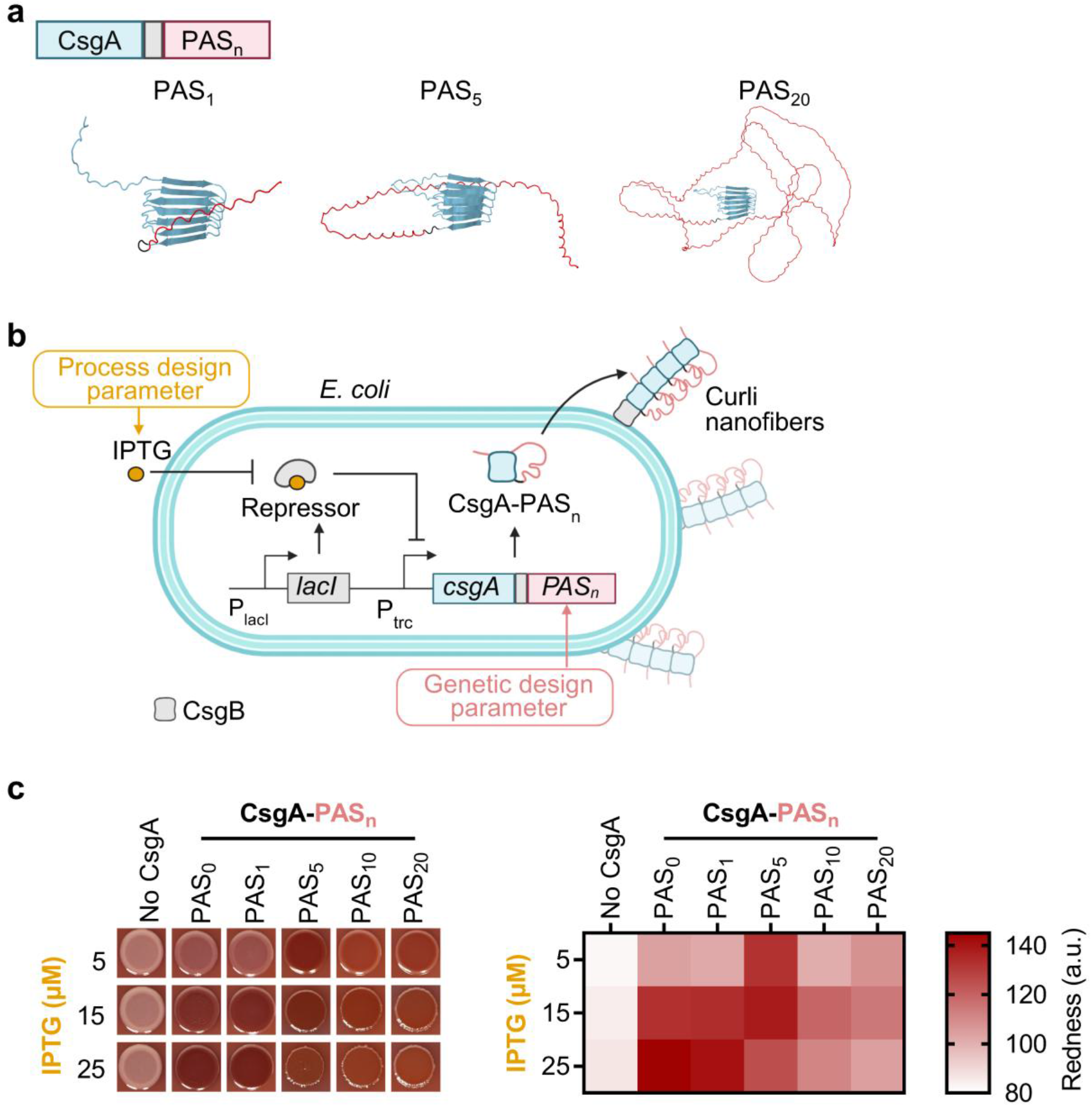
Design and construction of an engineered living hydrogel library. (**a**) Design of the building blocks. The fundamental building block consists of the CsgA protein (cyan) fused via a GSS linker (gray) to different numbers of repeats of the 20 amino acid PAS unit (1.7 kDa, red). Alphafold2-predicted^30,31^ structures for CsgA-PAS_n_ with n = 1, 5, and 20 are shown. (**b**) Schematic representation of the production strain for the engineered living hydrogel. *E. coli PHL628 ΔCsgA* bacteria were transformed with genetic constructs encoding for CsgA-PAS_n_ variants under the control of the IPTG-inducible P_trc_ promoter. As genetic and process design parameters, the number of PAS repeats as well as the concentration of IPTG were used, respectively. The CsgA-PAS_n_ proteins assemble the extracellular protein network forming the engineered living hydrogel. (**c**) Growth of a library of engineered living hydrogels. The different samples differ in number of PAS repeats and were grown in the presence of indicated IPTG concentrations. The hydrogels were grown on YESCA - Congo red agar plates for 48 h at 25 °C. (Left) Representative images of living hydrogels. (Right) Quantification of living hydrogel redness to estimate curli fibre production.

### 2.2 Characterization of the mechanical properties of the living hydrogel library

To characterize the mechanical properties of the living hydrogels by small amplitude oscillatory shear rheology, we grew the materials on a polycarbonate membrane with 0.4 µm pore size placed on YESCA agar which allowed both, diffusion of nutrients during growth phase (four days, 25 °C) and subsequent transfer of the living hydrogel to the rheometer (**Fig. 3a**). First, we performed an amplitude sweep and measured the storage (*G′*) and the loss modulus (*G″*) of CsgA-PAS_n_ living hydrogels grown in the presence of 15 µM IPTG and containing n = 0, 1, 5, 10 or 20 PAS repeats (**Fig. 3b-f**). At low strain amplitudes (0.02% - 0.08% effective strain), corresponding to the linear viscoelastic region (LVER), we observed *G′* being higher than *G″* indicating gel-like properties. Above 0.08% effective strain, *G′* progressively decreased, indicating yielding and disruption of the network structure. With increasing strain, the elastic component *G′* and the viscous component *G″* crossed between 1.4% and 3.2% effective strain, depending on the ELM variant. This behaviour is commonly observed in physically crosslinked hydrogels, where increasing deformation induces yielding and structural rearrangement^32,33^. Next, we determined *G′* and *G″* within the LVER (0.06% effective strain) for all hydrogel variants grown in the presence of different IPTG concentrations (5, 15, 25 µM) and containing different numbers of PAS repeats (**Fig. 3g**). *G′* showed a decreasing trend with increasing numbers of PAS repeats, likely due to an increase in flexible PAS polymers. Further, increasing IPTG concentrations led to higher *G′* values. Across all PAS_n_ and IPTG combinations, *G′* ranged from a minimum of 1227 ± 138 Pa (mean ± standard deviation, *n* = 3 living hydrogels) for CsgA-PAS_20_ and 5 µM IPTG to a maximum of 8615 ± 1196 Pa (mean ± standard deviation, *n* = 3 living hydrogels) for the CsgA-PAS_1_ variant grown at 25 µM IPTG. A two-way ANOVA analysis revealed that the genetic design parameter was the dominant source of variation for *G′*, accounting for 81.8% of the total variance (P < 0.0001, **Supplementary Table 3**). The process design parameter as well as the interaction among both also significantly affected *G′* accounting for 6.5% and 7.3% of total variance, respectively (P < 0.0001 and P = 0.0001, respectively, **Supplementary Table 3**).

**Figure 3.**
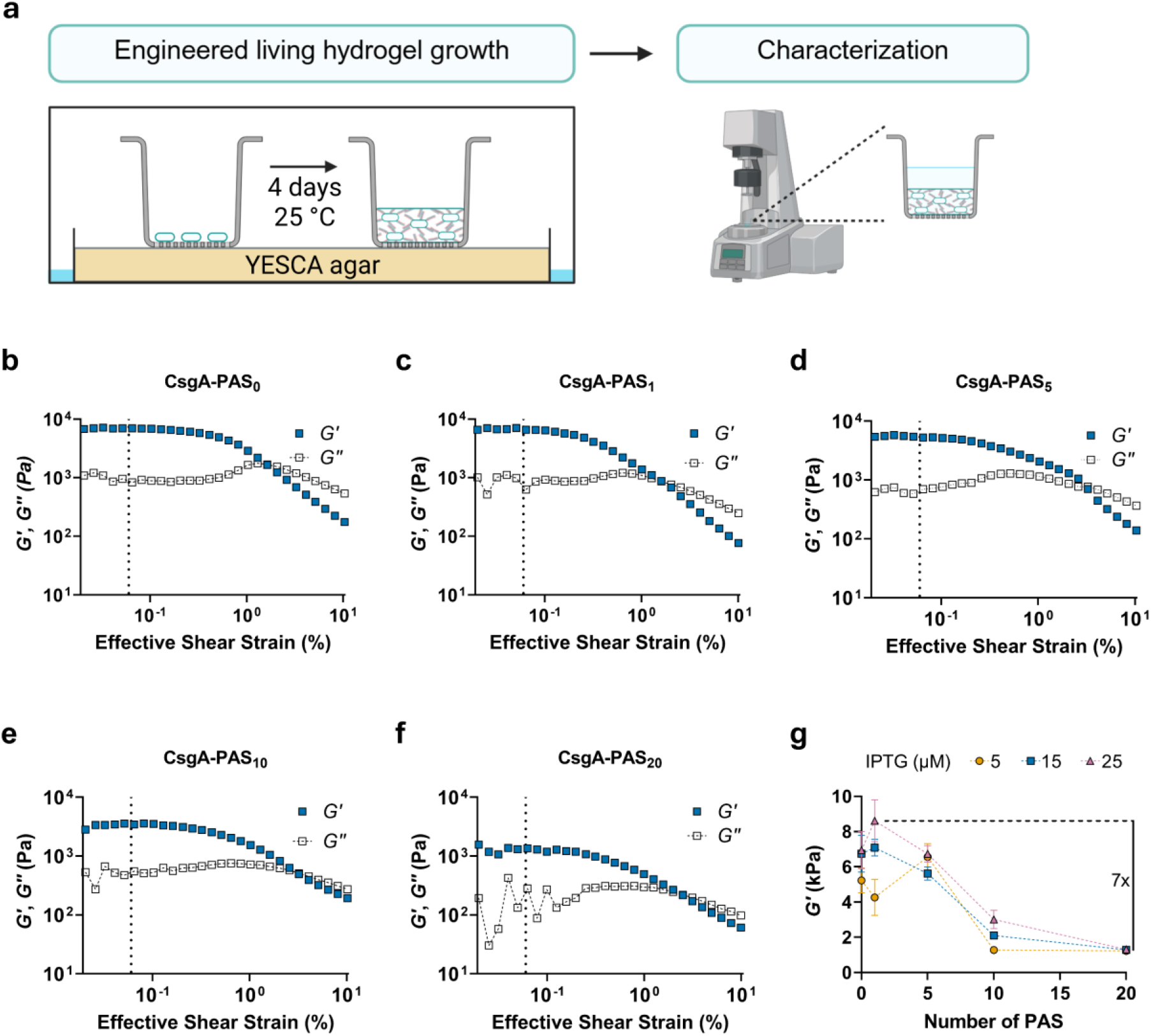
Determination of the mechanical properties of the engineered living hydrogel library. (**a**) Living hydrogel preparation. Living hydrogels were grown for four days at 25 °C in plate inserts placed on YESCA agar containing indicated IPTG concentrations prior to analysis by small-amplitude oscillatory shear rheology. (**b-f**) Oscillatory amplitude sweeps. The storage modulus (*G′*) and loss modulus (*G′′*) of living hydrogels containing indicated numbers of PAS repeats and grown in the presence of 15 µM IPTG were determined for 0.02-10 % effective strain at a frequency of 1 Hz. (**g**) Analysis of *G′*. *G′* was determined at 0.06 % effective strain at a frequency of 1 Hz for living hydrogels containing CsgA-PAS_n_ with n = 0, 1, 5, 10 or 20 and grown for four days in the presence of 5, 15, and 25 µM IPTG. Data points represent the mean and standard deviation for *n* = 3 living hydrogels prepared from the same culture batch and grown and characterized independently.

### 2.3 Evaluation of statistical models for *G′* prediction

We next evaluated the ability of different statistical models to predict *G′* from the genetic and process design parameters. We performed a comparative model selection study using linear regression, Gaussian process regression (GPR), and a Tabular Prior-data Fitted Network (TabPFN) (**Table 1**). These models were chosen to represent increasing levels of complexity, ranging from a simple parametric baseline, through a non-linear probabilistic, to a foundation-model-based approach. A log10-transformation was applied to *G′* values prior to fitting the Linear and GPR models due to the large dynamic range of measured *G′* values and the presence of non-constant variance across the response distribution. Log-transformation is commonly used to stabilize variance, reduce the influence of extreme observations^34,35^, and improve regression performance when response variables span multiple orders of magnitude^34,36^. Linear regression served as a baseline model, whereas GPR was implemented to capture nonlinear relationships and measurement noise. In contrast, TabPFN was trained directly on the original *G′* values. To ensure direct comparability between models, all predictions were converted back to the original scale, and all reported R^2^ and RMSE values were calculated using *G′* values in their original units (Pa).

**Table 1:** Predictive models tested to study the relationship between genetic and process design parameters and the resulting storage modulus (*G′*) of the living hydrogels.

| Model | Key parameters | Methodology |
| --- | --- | --- |
| Linear regression | Log10-transformed target | Baseline linear model trained on log-transformed $G'$ values using PAS repeat number and IPTG concentration as predictors<sup>41</sup>. |
| Gaussian process regression (GPR) | Matérn kernel ($\nu=1.5$); White Kernel | Non-linear probabilistic model trained on log-transformed $G'$ values, with the White Kernel accounting for experimental noise<sup>42</sup>. |
| TabPFN | Prior-Data Fitted Network (TabPFN) | Pretrained transformer-based model for tabular data trained directly on $G'$ values in the original scale, without target transformation or extensive task-specific preprocessing<sup>21</sup>. |

Implementing a robust validation strategy is critical when evaluating machine learning (ML) methods as it enables optimal model selection and provides a reliable estimate of its generalization to real-world data. To address this, rather than relying on simple random cross-validation across the dataset, model performance was assessed using two complementary validation strategies: Leave-One-Condition-Out (LOCO) cross-validation, in which individual PAS-IPTG combinations were withheld during training, and Leave-One-IPTG-Out (LOIO) cross-validation, in which an entire IPTG induction level was excluded from training and used for validation (see modelling methods section for technical details). LOCO evaluates the model’s ability to generalize across the experimental design space by predicting unseen PAS-IPTG combinations, while LOIO evaluates the models’ robustness to out-of-distribution data by holding out a complete IPTG concentration level during training^37^. In addition, the Ratio of Performance to Interquartile Distance (RPIQ) was calculated to assess prediction error relative to the variability of the measured data. RPIQ provides a normalized and dimensionless metric of predictive performance that facilitates comparison across models and properties^38,39^. Higher values of RPIQ indicate better predictive performance, with values of 1.7-2.0 generally considered indicative of a good model, 2.0-2.5 of a very good model, and above 2.5 of an excellent model^40^.

The three models were trained with *G′* values obtained for the CsgA-PAS_n_ living hydrogels with n = 0, 1, 5, 10 and 20 grown at 5, 15 and 25 µM IPTG. The prediction curves showed that both TabPFN and GPR captured nonlinear trends in *G′* more effectively than linear regression, particularly at 5 µM IPTG, where the linear model was unable to adequately reproduce the experimental observation (**Fig. 4a-c**). Across LOCO cross-validation, TabPFN demonstrated the strongest predictive performance with global R^2^ values of 0.713 and RMSE of 1098 Pa (**Table 2**). Additionally, when evaluating the error prediction over the measured data variability, TabPFN achieved the highest RPIQ (4.75), indicating that prediction errors were smaller relative to the interquartile spread of the *G′* measurements (**Table 2**). Under LOIO cross-validation, TabPFN produced slightly lower metric values than linear regression. The slight advantage of linear regression suggests that it has a greater ability to extrapolate to unknown IPTG concentrations when a full IPTG condition is not considered during training.

**Figure 4.**
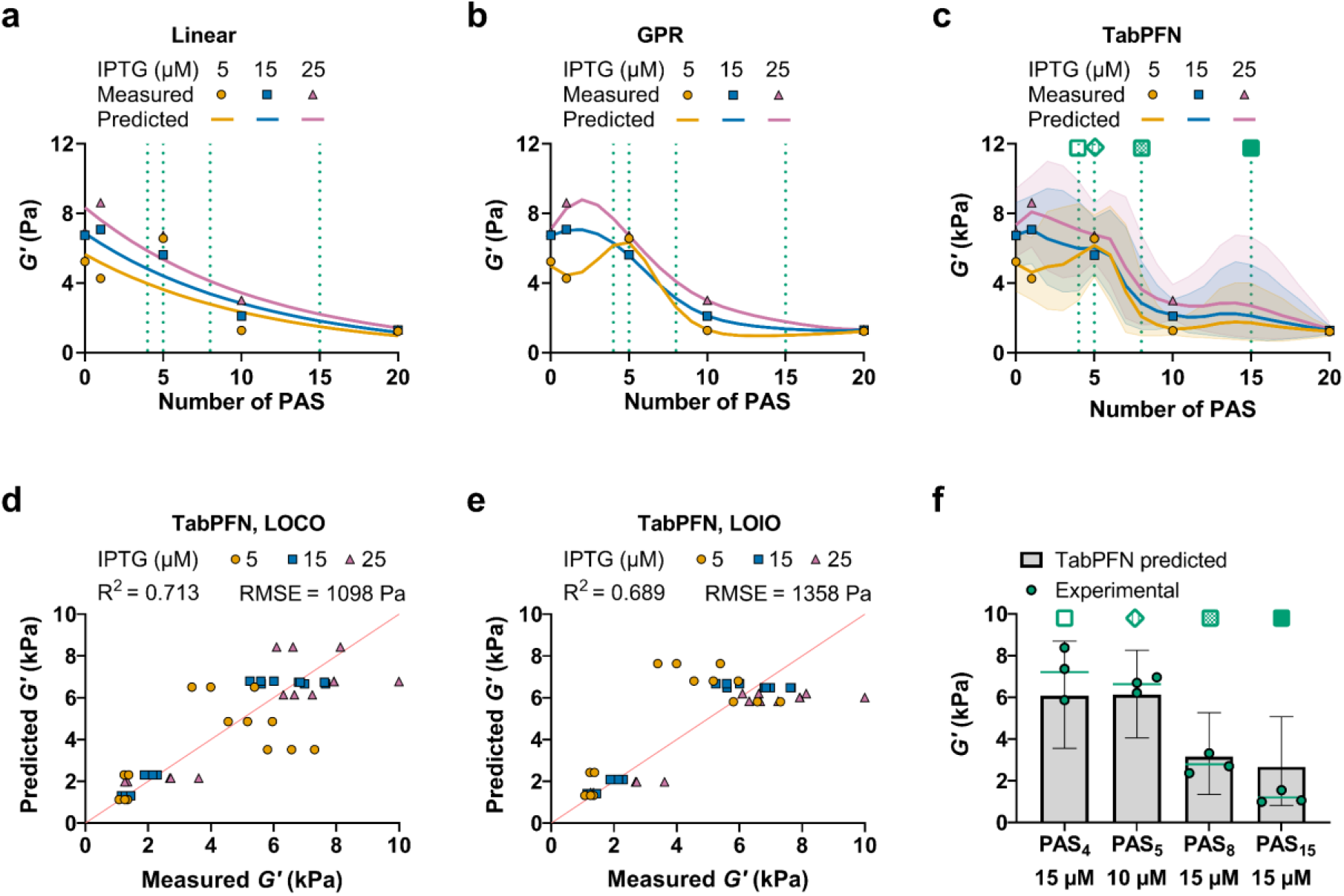
Training, cross-validation and experimental validation of predictive models for *G′*. (**a**) Linear regression, (**b**) Gaussian process regression (GPR) and (**c**) TabPFN models were trained with *G′* data of living hydrogels comprising CsgA-PAS_n_ with n = 0, 1, 5, 10 or 20 and grown in the presence of 5, 15, and 25 µM IPTG. In (**c**), the shaded areas represent the uncertainty predictions, corresponding to the 5^th^ and the 95^th^ percentiles of the predictive distribution. Solid lines represent the median of the predictive distribution. Marker shapes represent the mean of measured samples (*n* = 3 living hydrogels for each PAS-IPTG condition). Dash vertical lines and green symbols above curves correspond to experimental validation points. (**d, e**) Scatter plots of measured vs. predicted values by TabPFN resulting from (**d**) Leave-One-Condition-Out (LOCO) cross-validation, in which individual PAS-IPTG combinations were withheld during training, and (**e**) Leave-One-IPTG-Out (LOIO) cross-validation, in which an entire IPTG induction level was excluded from training and used for validation. The global R^2^ and Root Mean Square Error (RMSE) for the different cross-validation strategies are displayed. (**f**) Comparison between TabPFN-predicted and experimentally measured *G′* values for an independent validation dataset. Experimental validation of the model predictions was performed on unseen data from an independent experiment for CsgA-PAS_n_ hydrogels variants with indicated n and grown in the presence of indicated IPTG concentrations. Bars represent the predicted *G′* value, while error bars indicate the 90 % predictive interval (5^th^ - 95^th^ percentiles). Green dots represent individual experimental replicates of the test set, and the green horizontal line denotates the experimental mean (*n* = 3 living hydrogels for each PAS-IPTG condition). Green symbols above bars correspond to validation points shown in (**c**).

**Table 2.** Performance comparison of the Linear, Gaussian process regression (GPR), and TabPFN models for predicting *G′.* Performance was evaluated using Leave-One-Condition-Out (LOCO) and Leave-One-IPTG-Out (LOIO) cross-validation, as well as an independent experimental validation set. Model accuracy is reported using Root Mean Square Error (RMSE), R^2^ and the Ratio of Performance to Interquartile Distance (RPIQ). SD: standard deviation.

| Model | LOCO cross-validation |  |  | LOIO cross-validation |  |  | Independent experimental validation Set |  |  |
| --- | --- | --- | --- | --- | --- | --- | --- | --- | --- |
| | RMSE (mean ± SD) | R2 | RPIQ | RMSE (mean ± SD) | R2 | RPIQ | RMSE (mean ± SD) | R2 | RPIQ |
| Linear | 1189 ± 797 Pa | 0.692 | 4.39 | 1236 ± 286 Pa | 0.758 | 4.22 | 1917 Pa | 0.454 | 2.40 |
| GPR | 1405 ± 979 Pa | 0.559 | 3.71 | 2289 ± 634 Pa | 0.152 | 2.28 | 1121 Pa | 0.813 | 4.10 |
| TabPFN | 1098 ± 839 Pa | 0.713 | 4.75 | 1358 ± 475 Pa | 0.689 | 3.84 | 1001 Pa | 0.851 | 4.59 |

Furthermore, the measured dependence of *G′* on number of PAS repeats at 15 and 25 µM IPTG exhibits a shallow maximum followed by a gradual decline. In contrast, the 5 µM IPTG condition shows a more complex pattern comprising a local minimum at PAS_1_, a maximum at PAS_5_, and a subsequent decline (**Fig. 4a-c**). Consequently, when these characteristic data points are omitted during cross-validation, TabPFN is less able to reproduce these local features and instead predicts values that more closely follow trends observed in the remaining conditions, as reflected by the deviations observed in the measured versus predicted scatter plots (**Fig. 4d, e**). Under LOCO cross-validation, the largest deviations were observed for the PAS_1_ and PAS_5_ conditions at 5 µM IPTG, where *G′* was overestimated and underestimated, respectively (**Fig. 4d**). A similar effect is observed during LOIO validation when the entire 5 µM IPTG condition is excluded from training (**Fig. 4e**). Similar behaviour was observed for GPR (**Supplementary Fig. 2c, d**). The impact of excluding these observations was less apparent for linear regression because the model captured the overall trend but not the local maximum un minimum observed at 5 µM IPTG (**Supplementary Fig. 2a, b**), which may explain why linear regression performed better under LOIO.

Furthermore, we experimentally validated model predictions using a test set containing data from an independent experiment for previously unseen PAS repeats (n = 4, 8 and 15) at 15 µM IPTG and unseen IPTG level (10 µM) for the CsgA-PAS_5_ variant (**Supplementary Fig. 3**). Consistent with previous training results, TabPFN achieved the best performance on the independent test set, yielding the highest R^2^ value (R^2^ = 0.851) and the lowest prediction error (RMSE = 1001 Pa), outperforming both GPR (R^2^ = 0.813, RMSE = 1121 Pa) and linear regression (R^2^ = 0.454, RMSE = 1917 Pa) (**Table 2**). Similarly, on the independent test set, TabPFN achieved the highest RPIQ of 4.59, confirming robust predictive performance on previously unseen data (**Table 2**).

Although linear regression achieved slightly better predictive performance in LOIO cross-validation, TabPFN showed the strongest overall performance in the cross-validation schemes and the independent validation dataset. Our selection of TabPFN was not only guided by predictive accuracy but also by its fundamental architectural advantages. As a tabular foundation model, TabPFN benefits from extensive pretraining on diverse synthetic datasets^21^. This prior knowledge enables it to identify relationships in small datasets and make predictions from limited experimental observations. Unlike conventional ML models that require iterative parameter updating and fitting, TabPFN can be applied directly to datasets with minimal computational costs and manual effort associated with data preprocessing, model training, and hyperparameter selection^21^. These characteristics further motivated its selection for subsequent analyses.

Next, to assess the confidence associated with the predictions of the best-performing model, TabPFN, we calculated predictive uncertainty. TabPFN provides probabilistic predictions that enable uncertainty quantification alongside point estimates^21^. To visualize model predictions, we extracted the median prediction (50^th^ percentile) together with the 90% predictive interval (5^th^-95^th^ percentiles) (**Fig. 4c**). Importantly, in all experimental observations for the independent test set, *G′* fell within the predicted 90% predictive interval (**Fig. 4f, Supplementary Fig. 4a**). This indicates that the model provides realistic uncertainty estimates while maintaining good predictive accuracy. Collectively, these results demonstrate that TabPFN provides the strongest overall predictive performance and generalization capability for *G′* of engineered living hydrogels across previously unseen experimental conditions.

### 2.4 Extending the predictive model to other properties of the living hydrogels

Having demonstrated that data-driven methods can predict *G′* from genetic and process design parameters, we next investigated their capacity to predict additional properties. Specifically, we evaluated model performance for predicting the grown living hydrogel thickness, specific curli fibre content, and permeability to diffusion. These properties were selected because they represent key structural and functional characteristics of ELMs. Hydrogel thickness reflects the macroscopic material growth, whereas specific curli fibre provides information on biopolymer production and content. In contrast, permeability captures a distinct functional aspect of the material providing insight into how readily molecules can move through the living hydrogel matrix. This property is particularly relevant for drug-releasing living therapeutic materials^2^.

For thickness determination, we characterized the living hydrogels by optical coherence tomography (OCT, **Fig. 5a**, **Supplementary Fig. 5**). Specific curli fibre content was quantified by a Congo red spin-down binding assay, where curli content was normalized to cell density (**Fig. 5b**). Furthermore, we analysed the permeability of the living hydrogels to fluorescein isothiocyanate (FITC)-dextran (40 kDa) as a model macromolecule (**Fig. 5c**). Here, FITC-dextran solution was added to the upper compartment of a plate insert holding the living hydrogel, and diffusion into the lower compartment was quantified after 23 h.

**Figure 5.**
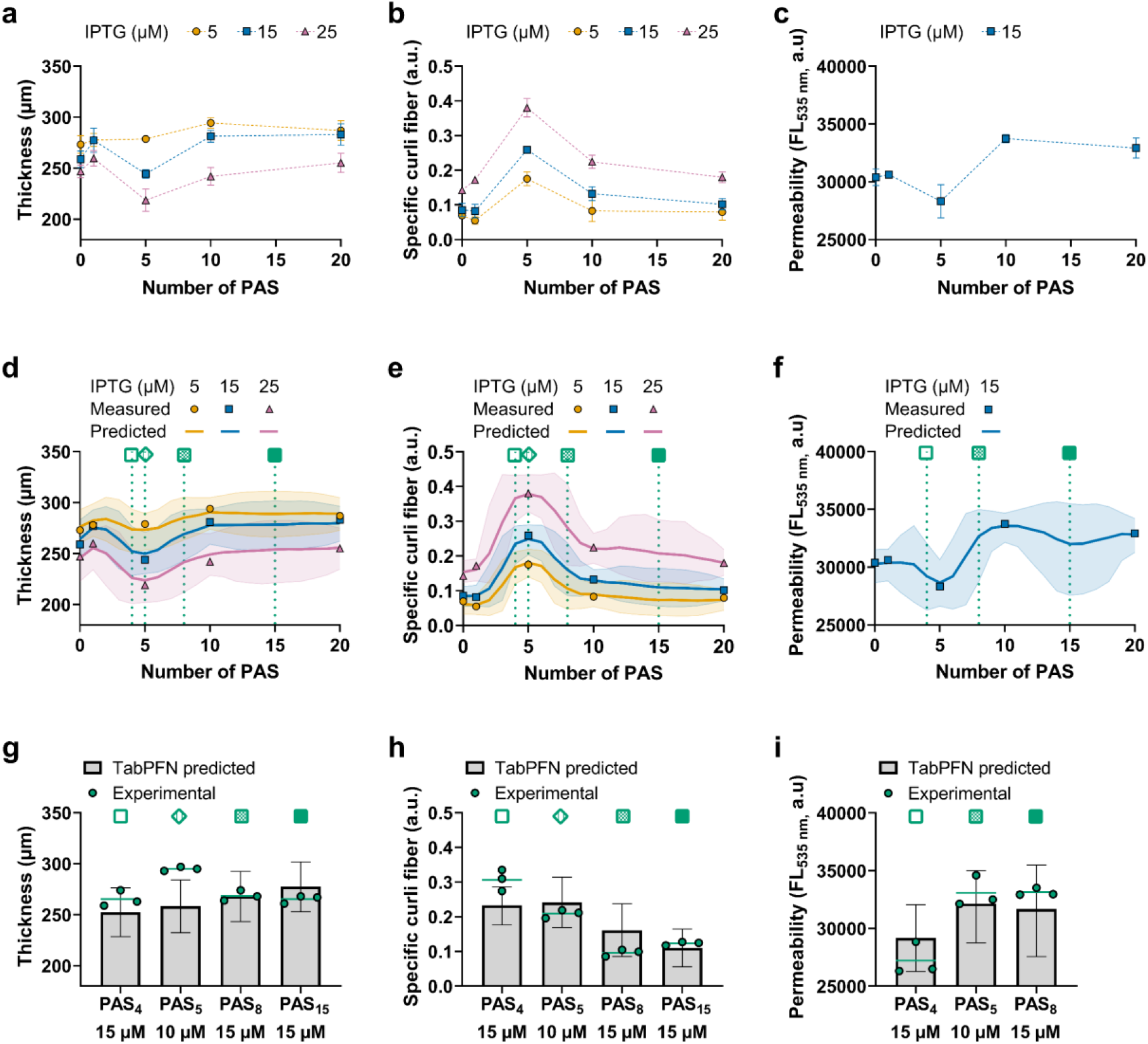
Extending the predictive model framework to further living hydrogel properties. Living hydrogel thickness (**a**), specific curli fibre content (**b**), and permeability to a 40 kDa FITC-dextran dye (**c**) was determined for variants containing CsgA-PAS_n_ with indicated n and grown in the presence of indicated IPTG concentrations. Data represent the mean and standard deviation (*n* = 3 living hydrogels for each PAS-IPTG condition). Using this data, TabPFN predictive models were implemented for (**d**) thickness (**e**) specific curli fibre content and (**f**) permeability. Solid lines represent the median of the predictive distribution. Marker shapes represent the mean of measured samples (*n* = 3 living hydrogels for each PAS-IPTG condition). The shaded areas represent the uncertainty predictions, corresponding to the 5^th^ and the 95^th^ percentiles of the predictive distribution. Dash vertical lines and green symbols above curves correspond to experimental validation points. (**g-i**) Comparison between TabPFN-predicted and experimentally measured values for the independent dataset. Experimental validation of the model predictions was performed on unseen data from independent experiments for CsgA-PAS_n_ hydrogels with indicated n and grown in the presence of indicated IPTG concentrations. Bars represent the predicted value, while error bars indicate the 90 % predictive interval (5^th^ - 95^th^ percentiles). Green dots represent individual experimental replicates of the test set, and the green horizontal line denotates the experimental mean (*n* = 3 living hydrogels for each PAS-IPTG condition). Green symbols above bars correspond to validation points shown in (**d-f**).

Analysis of variance revealed that PAS length was the primary determinant of both specific curli fibre and permeability. For curli fibre, PAS length accounted for 54.6% of the total variance, compared with 37.9% for IPTG concentration (**Supplementary Table 4)**. For permeability, PAS length explained 88.8% of the observed variance (**Supplementary Table 5**). In contrast, for thickness, IPTG concentration was the dominant factor, accounting for 54.0% of the total variance, whereas PAS length explained 24.3% (**Supplementary Table 6**).

Next, we trained TabPFN using thickness, specific curli fibre content, and permeability data for the CsgA-PAS_n_ living hydrogel variants with n = 0, 1, 5, 10, and 20. Model generalization was assessed using an independent test set comprising previously unseen CsgA-PAS_n_ variants with n = 4, 8, 15 at 15 µM IPTG; in addition, CsgA-PAS_5_ at unseen 10 µM IPTG was also used for thickness and specific curli fibre.

The model developed for specific curli fibre showed accurate predictive performance under LOCO, moderate under the more challenging LOIO evaluation, and a satisfactory predictive performance on the independent test set (R^2^ = 0.597, RPIQ = 2.25). The model for permeability showed good predictive ability on the independent test set (R^2^ = 0.690, RPIQ = 2.53) despite a negative R² during LOCO cross-validation (**Table 3, Supplementary Fig. 6**); this inconsistency is likely driven by the limited amount of data available for training and cross-validation, which makes R² and RPIQ to depend strongly on the variability present within each validation subset rather than reflecting a genuine loss of predictive accuracy. A similar pattern was observed for thickness, although the model yielded a negative R² and low RPIQ on the independent test set, the absolute prediction error remained relatively modest, with RMSE values ranging from 14 to 21 µm across all validation schemes, corresponding to approximately 5-8% of the overall mean thickness (267 µm) (**Table 3, Supplementary Fig. 6**). The degradation in R² and RPIQ is better explained by the narrower variability of the test set (SD = 13 µm) relative to the overall dataset (SD = 20 µm) than by a substantial loss of predictive accuracy, since R² is sensitive to the ratio between RMSE and the variance of the evaluation set itself. These results suggest that the decline in performance is largely influenced by the characteristics of the dataset rather than by a reduction of the model’s predictive capabilities. This observation highlights the importance of interpreting relative performance metrics (e.g., R² and RPIQ) alongside absolute error measures (e.g., RMSE) to provide a more complete assessment of predictive model performance, particularly for datasets exhibiting limited variability.

**Table 3.** Predictive performance of TabPFN models for hydrogel thickness, curli fibre content, and permeability. . Performance was evaluated using Leave-One-Condition-Out (LOCO) and Leave-One-IPTG-(LOIO) Out cross-validation, as well as an independent experimental validation set. Model accuracy is reported using Root Mean Square Error (RMSE), R^2^ and the Ratio of Performance to Interquartile Distance (RPIQ). SD: standard deviation.

| Property | LOCO cross-validation |  |  | LOIO cross-validation |  |  | Independent experimental validation Set |  |  |
| --- | --- | --- | --- | --- | --- | --- | --- | --- | --- |
| | RMSE (mean ± SD) | $R^2$ | RPIQ | RMSE (mean ± SD) | $R^2$ | RPIQ | RMSE | $R^2$ | RPIQ |
| Thickness | 14 ± 6 μm | 0.470 | 2.04 | 20 ± 7 μm | 0.038 | 1.48 | 21 μm | -1.505 | 0.71 |
| Specific curli fiber | 0.022 ± 0.008 a.u. | 0.927 | 4.27 | 0.053 ± 0.022 a.u. | 0.557 | 1.78 | 0.053 a.u. | 0.597 | 2.25 |
| Permeability | 1959 ± 1645 a.u. | -0.552 | 1.60 | — | — | — | 1629 a.u. | 0.690 | 2.53 |

Prediction curves, including the corresponding 90% uncertainty intervals were determined for all investigated properties (**Fig. 5d-f**). As previously observed for *G′*, some regions of the predictive distribution showed higher uncertainty, likely related to high data variability or limited data. The experimental observations for the independent test set for thickness, curli fibre amount and permeability fell within the predicted 90% predictive interval except for PAS_5_, 10 µM IPTG for thickness and PAS_4_, 15 µM IPTG for specific curli fibre that were marginally beyond the upper 90% boundary, by only 3.8% and 7.0%, respectively (**Fig. 5g-i**, **Supplementary Fig. 4b-d**). Incorporating additional values in regions of low predictive power, model performance could be improved.

We observed that permeability tended to decrease with increasing *G′* (Pearson r= 0.845, Spearman corelation ρ= 0.786, **Supplementary Fig. 7**), indicating that stiffer hydrogels exhibit lower macromolecular permeability. This inverse relationship is consistent with previously reported properties in synthetic hydrogels^43^.

Together, these results demonstrate that the modelling framework can be extended beyond mechanical properties to predict structural and functional characteristics of engineered living hydrogels from small datasets.

### 2.5 Prediction of design parameters for living hydrogels with desired properties

We finally evaluated whether we could use TabPFN to predict genetic and process design parameters that yield living hydrogels with desired properties. Typically, inverse design approaches are performed for this purpose^44,45^, however the inverse function of TabPFN cannot be computed analytically. We therefore approached this task through systematic exploration of the predicted design space rather than direct inverse modelling. This approach is feasible because of the low dimensionality of the design space. Although leveraging TabPFN’s predictive capabilities within a Bayesian Optimization^46^ pipeline represents an attractive direction for inverse design, such extensions are likely to be more beneficial for exploring larger or higher dimensional design spaces, and were therefore beyond the scope of the current study. Using the previously implemented predictive models we built the possible design spaces, and we use the independent test-set to study the behaviour of the predictions. To generate the predictive spaces, we performed a 2D parameter sweep of PAS repeats and IPTG concentrations, and we generated independent predictive spaces for *G′*, thickness, specific curli fibre, and permeability using separate TabPFN models (**Fig. 6**). These predictive spaces were queried to identify design parameter combinations predicted to yield the closest value to a target property.

**Figure 6.**
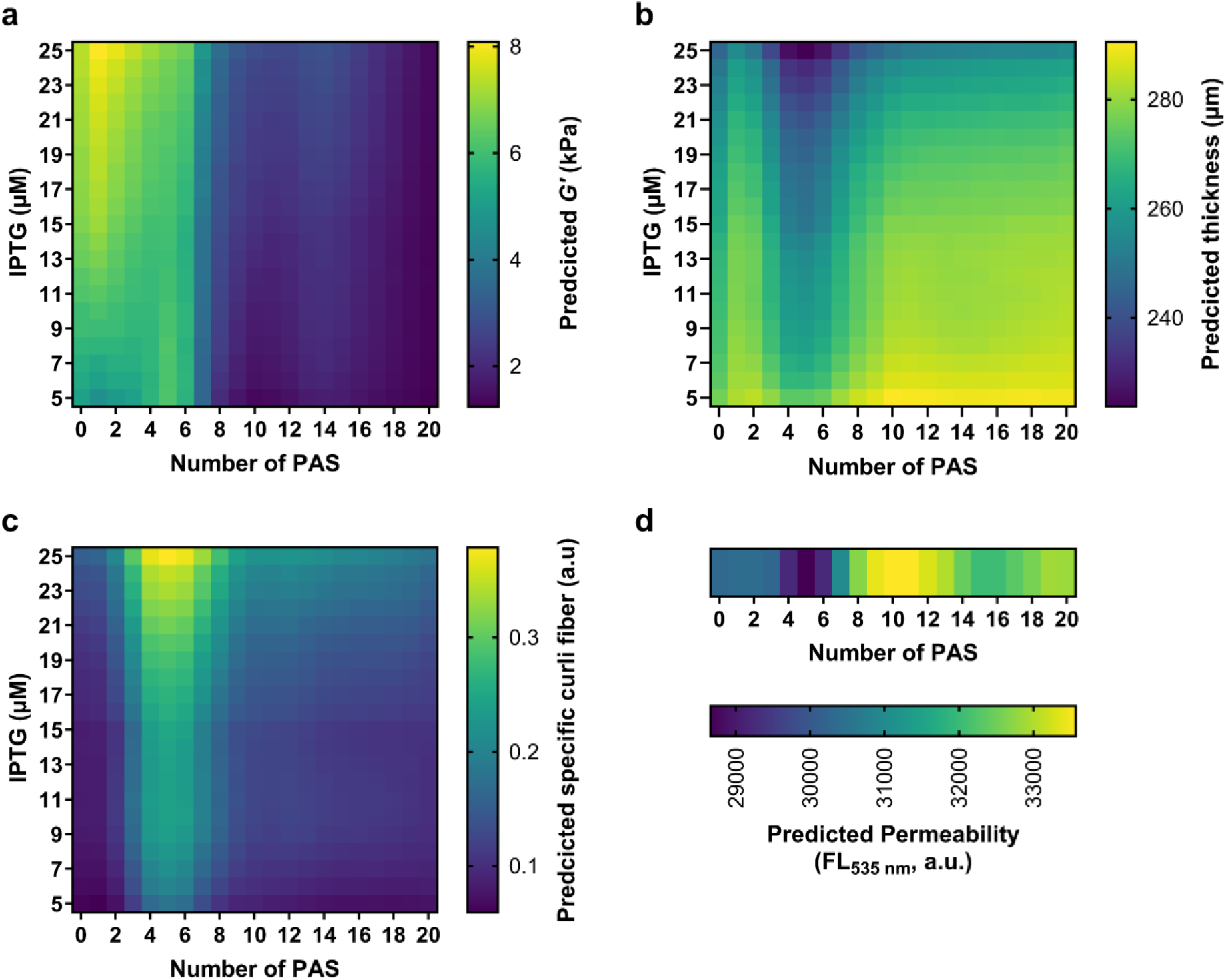
Implementation of a property-guided prediction approach to determine design parameters that yield living hydrogels with desired properties. Using the trained and validated TabPFN models, heatmaps of the design parameter space defined by the number of PAS repeats and the IPTG concentration were implemented for (**a**) *G′*, (**b**) thickness, (**c**) specific curli fibre content and (**d**) permeability. The colour scale represents the model-predicted properties of the living hydrogel. These predictive spaces can be queried to identify design parameter combinations predicted to yield living hydrogels with desired properties.

For example, when targeting a living hydrogel with a *G′* of approximately 2800 Pa, corresponding to the mean of the PAS_8_, 15 µM IPTG unseen condition, the model identified several parameters combinations predicted to yield a similar *G′*. Notably the original experimental design parameter combination was only ranked in the 10^th^ position. This is expected because the mapping between design parameters and material properties is not unique, multiple genetic and process design parameter combinations can produce comparable hydrogels phenotypes. Similar behaviour was also observed for all the other independent, unseen conditions (**Supplementary Table 7**), highlighting that the framework identifies feasible solutions rather than attempting to recover a single exact design.

To address this inherent redundancy, we extended the search from a single property to a multi-property objective incorporating *G′*, thickness, and specific curli fiber simultaneously. In this approach, candidate designs are identified based on proximity to a target material property within a normalized three-dimensional property space. Considering multiple properties substantially improved recovery of experimentally relevant design parameters across all unseen conditions (**Supplementary Table 8**). For instance, the distance (Manhattan distance, see methods) between the predicted and experimental design parameters for the PAS_8_, 15 μM IPTG condition decreased from 14 to 1, demonstrating that richer phenotypic descriptions more effectively constrain the design space.

Ultimately, we showed that user preferences provide an additional layer of selection by incorporating practical design constraints, such as favouring lower IPTG concentrations (for reduced cellular stress) or restricting PAS_n_ selection to available cloned variants. Combined with multi-property performance criteria, these preferences can guide the optimal selection of design parameters that meet both the functional requirements and the experimental constraints (**Supplementary Table 9**).

This approach demonstrated how predictive models can support property-guided design of engineered living hydrogels. By prioritizing a small number of candidate genetic and process design parameters, the framework can reduce experimental search efforts and further accelerate the rational design of living hydrogels with desired characteristics.

To provide an improved design space framework for future work, we retrained TabPFN with the hitherto obtained complete data set for *G′*, thickness, curli fibre and permeability including the samples previously reserved for independent testing. After retraining, the prediction uncertainty decreased, as reflected in narrower uncertainty intervals particular in regions with previously limited data (**Supplementary Fig. 8**). Model performance was generally maintained across properties, with the largest improvement observed for permeability, as evidence by a 45% increase in R^2^ and 50 % reduction in RMSE (**Supplementary Table 10)**. To make the models more accessible and easier to use, the final models trained on the entire dataset, were integrated into an interactive web application developed using the Streamlit platform^47^ (**Supplementary Fig. 9**). This tool allows to quickly explore the design space and provides an intuitive interface for predicting both material properties and candidate design parameters, without requiring programming skills.

## 3. Discussion

A central challenge in the ELM field is establishing quantitative relationships between programmable biological design and process parameters and resulting material properties and functions. Here, we demonstrate that the storage modulus (*G′*), curli fibre content, permeability and thickness of curli-based living hydrogels can be predicted directly from two experimentally accessible design parameters: (i) the number of genetically encoded biopolymer length (genetic parameter) and (ii) induction level (process parameter). Using a small experimental dataset, TabPFN models precisely predicted these properties and generalized to independent, previously unseen conditions (57 points in total, 45 for training and 12 for independent validation). Furthermore, we developed an inverse-design strategy based on the TabPFN models to identify candidate design parameter for growing living hydrogels with desired material properties. Together, these results establish a quantitative framework for the forward and inverse design of living hydrogel properties directly from programmable design parameters.

Our results identified the genetic design parameter as the primary variability determinant of *G′*, curli fibre production, and permeability. PAS polypeptides emulate key biophysical properties of polyethylene glycol (PEG), including high hydrophilicity, large hydrodynamic volume, and neutral charge^26,27^, which may influence multiple stages of curli biogenesis and consequently alter living hydrogel properties. First, PAS polypeptides may impair secretion through the CsgF-CsgG complex, particularly long PAS repeats, since the pore diameter of this secretion complex is approximately 1 nm^25^. Second, PAS polypeptides may interfere with folding and self-assembly of CsgA, by introducing steric constraints at its C-terminus, potentially disrupting intermolecular interactions required for efficient curli fibre formation^48^. Consistent with these interpretations, it has been previously shown that fusing peptides to CsgA can reduce its secretion efficiency leading to lower curli fibre formation, as reported in the pioneer BIND-CsgA platform, where large fusion proteins can result in decreased curli formation^11^. In addition, previous studies have shown that fusing different peptides to CsgA can change the mechanical properties of CsgA-based hydrogels. For example, trefoil factor (TFF) fusions yielded hydrogels with distinct *G′* values, which were attributed to differences in aggregation behaviour and interfibre interactions introduced by the fused domains^17^. A second layer of control is provided by the process design parameter, which was the dominant factor explaining variability in hydrogel thickness. The effects of IPTG likely arise from changes in CsgA-PAS_n_ expression levels as well as potential cellular stress associated with overexpression. Consequently, different combinations of PAS repeat length and IPTG concentration may generate matrices with different compositions and structural organizations, yielding living hydrogels with different properties. Overall, these findings establish a programmable living hydrogel design platform in which material properties can be genetically encoded and further tuned by process-based parameters.

Extracting meaningful relationships from small tabular datasets, typical of the ELM field, requires suitable modelling approaches. Among the models evaluated, the foundation model TabPFN, achieved the highest predictive accuracy for *G′*, particularly for unseen genetic and process parameters. The model effectively captured non-linear relationships while providing well-calibrated uncertainty estimates in a zero-shot setting, meaning that predictions could be generated without explicit training or fine-tuning on the target dataset. Despite its recent introduction (2025^21^), TabPFN has already demonstrated promise across diverse applications where data is limited, including the guided design of wood composites^22^ and clinical diagnostic and prognostic prediction tasks^23,24^. Notably, these studies used datasets containing approximately 200-1,500 datapoints, substantially larger than the 57 datapoints used here (45 for training and 12 for independent validation). Although the primary focus of this study was the prediction of hydrogel mechanics, the framework also proved applicable to additional material properties including curli fibre content, macromolecular permeability and thickness. Collectively, these results highlight the potential of transformer-based foundational models for predicting ELM properties from small datasets.

A key objective in ELMs, and in materials science more broadly, is to identify design parameters required to achieve a desired material property. Our property-guided design framework based on a grid-based strategy addresses this challenge. While this approach is suitable for small design spaces, it may become impractical as the dimensionality and size of the design space increase; in such cases, more sophisticated methods, such as Bayesian optimisation^46^, may be more appropriate. Nevertheless, experimental datasets in the ELM field often contain only tens of measurements, making the approach highly relevant in practice. By narrowing the experimental search space to a limited number of promising candidates, this approach paves the way for property-driven design of engineered living hydrogels.

Despite the promising predictive performance achieved in this study, several opportunities for improvement remain. The predictive models were trained with a small dataset; additional data would likely improve prediction accuracy and reduce uncertainty. The design space was limited to two design parameters, one genetic and one process, and as with most predictive models, extrapolation beyond the design space is likely to be less reliable. Although these were sufficient to accurately predict several material properties, incorporating additional genetic and process parameters could broaden the design space as well as increase the range of predictable properties. Our analysis focused on predicting material properties directly from design parameters because these inputs are readily accessible and available prior to material prediction.

Future work could investigate whether incorporating measured material properties into model training further improves prediction performance. For example, structural characteristics such as curli fibre content or hydrogel thickness may provide additional information that enhances the prediction of properties such as *G′* or permeability. Important to note that while this may enhance accuracy, it would reduce one of the key advantages of the current framework by requiring previous knowledge of some material properties, which might not be accessible or require additional experimental work. An important advantage of data-driven models is their ability to identify relationships between design parameters and material properties even when the underlying biological mechanisms are not fully understood. This is particularly valuable in our system, where PAS repeats fused to CsgA and IPTG concentrations may influence multiple biological processes of curli biogenesis, including secretion, folding, and assembly. However, they do not provide a mechanistic understanding of the processes driving changes in material properties, which could be beneficial in uncovering new layers of rational design of this platform. Addressing these limitations will further improve and expand the capabilities of data-driven approaches for living hydrogels design.

Overall, our results demonstrate that predictive models trained on limited experimental datasets can accurately capture relationships between programmable genetic and process design parameters and resulting material properties. Here, we establish a quantitative framework for the predictive and property-guided design of engineered living hydrogels. We anticipate that this approach will facilitate the transition from empirical optimisation to predictive, property-driven design, ultimately accelerating the rational development of ELMs with tailored mechanical, structural, and functional characteristics.

## 4. Methods

### 4.1 Experimental methods

#### Plasmids

The pBbE1a-CsgA plasmid (kindly provided by Prof. Neel S. Joshi, Northeastern University, USA)^11^ was used to clone the CsgA-PAS_n_ encoding sequences. To build the CsgA variants containing different number of repeats of the 20-amino acid core PAS polypeptide (ASPAAPAPASPAAPAPSAPA)^26^ at the C-terminus of CsgA (CsgA-PAS_n_), we employed a cloning strategy similar to that described by Schlapschy et al.^26^ Initial PAS sequences were assembled in a vector surrounded by *Sap*I restriction sites. Then the initial PAS cassettes were extracted by *Sap*I digestion and introduced at the C-terminus of CsgA via a *Sap*I site. The resulting pCsgA-*Sap*I-PAS_n_ vectors can then be opened by *Sap*I restriction digestion and additional PAS cassettes can be ligated to generate larger PAS repeats at the C-terminus of the CsgA. All the sequences of the open reading frames were verified. Additional information on the DNA sequences encoding the CsgA-PAS_n_ variants is provided in **Supplementary Table 1** and **2.**

#### 3D protein structure

3D protein structure prediction of CsgA-PAS_n_ variants was generated using ColabFold^31^, employing the AlphaFold2-pTM model^30^. Five models were generated with three cycles and ranked according to pLDDT scores. The top ranked models for each CsgA-PAS_n_ were selected for visualization and processing in Jmol v16.1.1^49^.

#### Characterization of biofilms on YESCA - Congo Red (CR) agar plates

Congo Red (CR) staining on agar plates was adapted from previous protocols^11,50^. CR (Sigma-Aldrich, Germany) and Brilliant Blue G250 (Sigma-Aldrich) were prepared at a stock concentration of 10 mg mL^-1^ in ultrapure water, filter sterilized and stored at 4 °C. YESCA agar plates (10 g L^-1^ casamino acids, 1 g L^-1^ yeast extract, 20 g L^-1^ agar, Carl Roth, Germany) were supplemented with 100 µg mL^-1^ ampicillin, indicated concentrations of IPTG (from a 0.1 mM stock solution in ultrapure water, Carl Roth), 50 µg mL^-1^ CR and 10 µg mL^-1^ Brilliant Blue G250. 5 µL of overnight liquid cultures were spotted on these plates and CR staining was observed after incubation for 48 h at 25 °C.

Images of agar plates were acquired using a Canon EOS R50 camera equipped with an RF-S18-45 mm F4.5-6.3 IS STM lens. CR staining was quantified using a custom macro implemented in Fiji (ImageJ v1.54i)^51^ including the BioVoxxel toolbox v2.6^52^. Living hydrogel regions of interest (ROIs) were identified by image segmentation from 8-bit grayscale images following contrast enhancement (saturated=0.01% normalize), variance filtering (radius = 15 pixels), automatic thresholding with Li algorithm, and morphological processing (10 dilations and 15 closing operations). The image area containing the living hydrogels spots was manually selected to exclude background artifacts originating from the edge of the Petri dish. Binary masks were further refined by hole filling, erosion (80 iterations), and opening (45 iterations). The resulting ROIs were transferred to the brightness channel of an HSB (Hue, Saturation, Brightness) stack generated from the original image, where mean pixel intensity values were measured. These measurements were used as a quantitative estimation of the curli fibre amount produced by the engineered living hydrogels.

#### Living hydrogel production in permeable plate inserts

The engineered living hydrogels produced by *E. coli* PHL628 *ΔCsgA* transformed with the plasmids encoding the CsgA-PAS_n_ variants were grown from a glycerol stock overnight in LB Medium (10 g L^-1^ tryptone, 5 g L^-1^ yeast extract, 10 g L^-1^ NaCl, Carl Roth) containing 100 µg mL^-1^ ampicillin at 25 °C and 150 rpm. OD_600nm_ was measured, LB medium removed by centrifugation at 3200 x g and OD_600nm_ adjusted to 20 in YESCA liquid medium (10 g L^-1^ casamino acids, 1 g L^-1^ yeast extract, Carl Roth). Cells were subsequently added to permeable plate inserts containing a 0.4 µm pore-size polycarbonate membrane, previously placed on top of a YESCA agar plate supplemented with 100 µg mL^-1^ ampicillin and indicated concentrations of IPTG. They were placed inside a polypropylene plastic box (550 mL capacity, Emsa, Germany) containing a tissue wetted with 10 mL of water and incubated for four days at 25 °C to prevent drying out. For mechanical characterization, ELMs were grown in 12 mm plate inserts (VWR, USA, European cat. No. 734-2731), whereas for permeability measurements, 6.5 mm inserts (Transwell®, Corning, USA, cat. No. 3413) were used.

#### Rheological measurements

The storage modulus (*G′*) of engineered living hydrogels was measured by small-amplitude oscillatory shear (SAOS) rheology using an MCR302e rheometer (Anton Paar, Austria) equipped with a parallel-plate geometry (PP08/S and P-PTD200, Anton Paar). Data acquisition was performed using the RheoCompass Software v1.32. To measure living hydrogels in native conditions, plate inserts (VWR, USA, European cat. No. 734-2731) containing the ELMs were mounted on the bottom plate of the rheometer and using a double-sided adhesive film. To minimize dehydration during testing, 300 μL of phosphate-buffered saline (PBS) was added to the insert. Measurements were performed at 25 °C under a normal force of 10 mN. Amplitude sweeps were conducted at a constant frequency of 1 Hz over an effective strain deformation of 0.02-10% to identify the linear viscoelastic region (LVER). Subsequently, *G′* was measured at a frequency of 1 Hz and an effective strain of 0.06% within LVER. The frequency was selected based on previous frequency-sweep analyses of related curli-based hydrogels reported by our group and others^53,17^. To calculate the effective strain applied to the living hydrogels, the gap associated with the plate insert and adhesive layer was measured using the rheometer under the same experimental conditions as those used for living hydrogel characterization, including a normal force of 10 mN. The combined height of the insert and adhesive layer was determined to be 252 ± 1 μm (mean and standard deviation for three different plate inserts). Considering this additional 252 µm and the total sample gap (living hydrogel, insert and adhesive layer), provided also by the rheometer, the effective strain applied to the ELMs was calculated as 0.06%. For each sample, five consecutive measurements were recorded and averaged to obtain the final *G′* reported value. For each PAS-IPTG condition *n* = 3 living hydrogels prepared from the same culture batch were grown and characterized independently.

The formula used to calculate the effective shear strain was derived from *Shear strain* = Δx/L, where *Δx* is the displacement and L the height. Considering *Δx* constant it resulted:

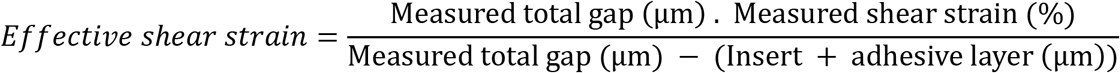

#### Determination of living hydrogel thickness

Engineered living hydrogels grown on plate inserts were imaged using a Ganymede GAN312 OCT (Thorlabs Ganymede, USA) with a central wavelength beam of 880 nm and a Thorlabs LSM03 scanning objective. Cross-sectional images were acquired using the ThorImage OCT v5.6 software with the following parameters: an A- and B-scan averaging of 8; an image frequency of 40 kHz; and the Hann Apodization window. A length of 11.8 mm was imaged. The thickness for each sample was calculated by dividing the area of the living hydrogel layer by its length. The refractive index required for the thickness calculation was set to 1.4, as determined for similar curli-based ELMs in our previous study^53^. For each PAS-IPTG condition *n* = 3 living hydrogels prepared from the same culture batch were grown and characterized independently.

The area of the OCT cross-sectional images was determined in Fiji (ImageJ v1.54i)^51^ including the BioVoxxel toolbox v2.6^52^ with a custom macro. Images were converted to 8-bit grayscale and pre-processed using a median filter (radius = 10 pixels). Background signal was corrected using the Convoluted Background Subtraction plugin (rolling=250 pixels) and the image contrast was enhanced (saturated = 0.01% normalize). ROIs were segmented by automatic thresholding using the Li algorithm. Binary masks were subsequently refined through morphological operations, including closing (35 iterations) and erosion (5 iterations). Objects were then detected using the Analyse Particles function, and the area of the ROIs were measured automatically.

#### Spin-Down quantitative Congo Red (CR) binding assay

The protocol was adapted from previously described studies^53,54^. ELMs grown on plate inserts were resuspended in PBS and pelleted at 6000 x g for 10 min. The pellet was resuspended in 0.075 mM CR (Sigma-Aldrich) in PBS and incubated at room temperature for 10 min. The mixture was then centrifuged at 20,000 x g for 10 min at 4 °C and the absorbance of CR in the supernatant was measure at 490 nm in a Tecan Spark multimode microplate reader (Tecan Trading AG, Switzerland). Specific curli fibre production was calculated by subtracting the absorbance obtained for the 0.075 mM CR solution from that of the ELMs and normalized by the OD_600nm_ of the resuspended ELM. For each PAS-IPTG condition *n* = 3 living hydrogels prepared from the same culture batch were grown and characterized independently.

#### Living hydrogel permeability

Technical details for this assay were adapted from previous studies^55,56^. Living hydrogels grown in 6.5 mm plate inserts with a 0.4 µm pore-size polycarbonate membrane as above described (Transwell®, Corning, USA, cat. No. 3413) were placed in black 24-well microplate Sensoplates (Greiner, Germany, cat. No. 662892). Samples were preincubated for 1 h at 25 °C in PBS, with 600 µL added to the plate well and 75 µL to the insert. Then, 25 µL of freshly prepared fluorescein isothiocyanate (FITC)-Dextran 40 kDa at 2 mg mL^-1^ was added to the insert to achieve a final concentration of 0.5 mg mL^-1^. Samples were incubated for 23 h in the dark at 25 °C inside a polypropylene plastic box (1.9 L capacity, LocknLock, Germany) containing a tissue wetted with 40 mL of water. After incubation, the plate inserts were carefully removed, and the fluorescence intensity of the Sensoplate wells were measured with an excitation wavelength of 485 nm and an emission of 535 nm in a Tecan Spark multimode microplate reader (Tecan Trading AG). An insert containing PBS only was used as a blank. For each PAS condition *n* = 3 living hydrogels prepared from the same culture batch were grown and characterized independently.

### 4.2 Modelling methods

#### Data Preprocessing

To address the broad dynamic range of the measured storage modulus *G′* (1,075 Pa-9,996 Pa) and the associated heteroscedasticity (experimental variance scaled proportionally with *G′* magnitude), target values were log-transformed:

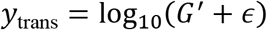

where *ε* = 10^−6^ Pa is a small numerical stabilizer to prevent mathematical singularity at zero. All predictions generated by log-domain models were transformed back to the physical scale (*G*^′^ = 10^ŷtrans^) prior to statistical evaluation and metric calculation.

A log transformation of the target was applied prior to fitting the linear regression and Gaussian process regression (GPR) models, which improved predictive performance for both. The same transformation was also evaluated for TabPFN but degraded its performance, and it was therefore not applied in the final TabPFN models. This is consistent with TabPFN’s regression approach, which predicts a discretized distribution over the target space rather than optimizing continuous squared error^21^.

For Linear regression and GPR, predictor variables (*N*_PASn_ and *C*_IPTG_) were standardized using *z*-score normalization to achieve zero mean and unit variance across the training set:

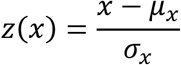

where *μ_x_* and *σ_x_* denote the empirical mean and standard deviation of feature *x*, respectively. For TabPFN, predictor variables were left unscaled on their original numerical scale, as the model performs internal feature normalization automatically and does not require manual preprocessing^21^.

#### Linear Regression

Ordinary Least Squares (OLS) multiple linear regression was implemented operating in the log-transformed target space. The model specifies a linear relationship between the standardized predictors and the logarithmic storage modulus:

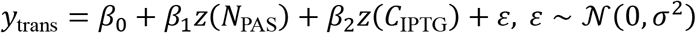

where *β*_0_ is the intercept, *β*_1_ and *β*_2_ are the partial regression coefficients corresponding to PAS length and IPTG induction, and *ε* represents normally distributed residual error. Parameter estimation was conducted by minimizing the sum of squared residuals:

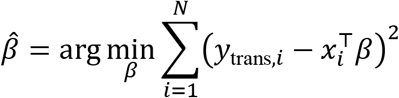

#### Gaussian Process Regression (GPR)

GPR was implemented using scikit-learn (v1.3). GPR models the target distribution as a non-parametric Bayesian prior over functions:

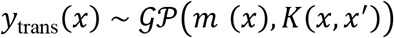

where *m*(*x*) = 0 is the prior mean function, and *K*(*x*, *x*^′^) is a composite covariance kernel defined as:

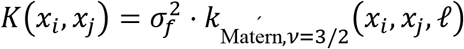

The signal component utilizes a Matérn ^3^ kernel to capture smooth non-linear interactions across the predictor space:

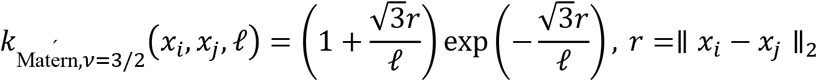

where ℓ is the characteristic length-scale (initial value ℓ = 1.0) and *σ^2^_f_* is the constant signal variance scale (bounded in [10^−3^, 10^3^]). The total covariance matrix of noisy target observations *y*_trans_ is given by: *K_y_* = *K*(*X*, *X*) + *σ^2^_n_I_N_*, where *σ^2^_n_* ∈ [10^−6^, 10^1^] models independent additive experimental noise via the identity matrix *I_N_*. Kernel hyperparameters *θ* = {*σ^2^_f_, l, σ^2^_n_*} were optimized via maximum likelihood estimation (MLE) by maximizing the log-marginal-likelihood (LML) of the training observations:

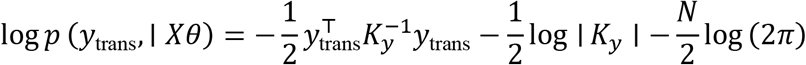

Optimization was executed using the L-BFGS-B algorithm with 10 random optimizer restarts (n_restarts_optimizer=10) to prevent convergence to local minima. Target normalization was applied prior to kernel evaluation (normalize_y=True).

#### TabPFN (Prior-Data Fitted Network)

TabPFN was employed as a foundation model for tabular data^21^. The model is based on a Transformer architecture that was pretrained on millions of synthetic tabular datasets generated from structural prior distributions, enabling strong predictive performance on small datasets without task-specific training, parameter updates, or hyperparameter optimization.

Rather than being trained on the experimental data in the conventional sense, TabPFN performs inference through in-context learning: a single forward pass in which the training examples (features and corresponding target values) and the test examples (features only) are jointly provided as input to the pretrained network. The model uses its attention mechanism to relate the test queries to the labelled training context, implicitly approximating a Bayesian posterior predictive distribution over the target conditioned on the provided examples^21^. No gradient-based parameter updates are performed on the training or test data at any stage; the same frozen, pretrained weights are reused for every fold, with the fold-specific training and test partitions supplied only as input at inference time.

#### Ratio of Performance to Interquartile Distance (RPIQ) Metric

RPIQ provides a normalized and dimensionless metric of predictive performance that facilitates comparison across models and properties^38,39^.

The RPIQ was calculated as:

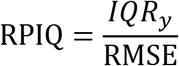

Where RMSE is the root mean square error of the model predictions, and *IQR_y_* is the interquartile range of the observed stiffness values calculated as:

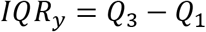

with *Q*_1_ and *Q*_3_ representing the first and third quartiles of the observed values, respectively. Higher values of RPIQ indicate better predictive performance, with values of 1.7-2.0 generally considered indicative of a good model, 2.0-2.5 of a very good model, and above 2.5 of an excellent model^40^.

A limitation of RPIQ is that it is normalized by the interquartile range of the test set itself rather than the full training dynamic range. Consequently, for test sets in which the data spans a narrow range of observed values, RPIQ can appear low even when the absolute prediction error is small relative to the overall training range. This effect is most pronounced for test sets with small sample sizes, where the interquartile range is estimated from few points and is therefore less stable.

#### Model Validation and Evaluation

The models were trained with values obtained for the CsgA-PAS_n_ living hydrogels with n = 0, 1, 5, 10 and 20 grown at 5, 15 and 25 µM IPTG (45 measurements, for each of the 15 PAS-IPTG conditions *n* = 3 living hydrogels prepared from the same culture batch and grown and characterized independently). Models were cross-validated using two complementary methods, Leave-One-Condition-Out (LOCO) and Leave-One-IPTG-Out (LOIO) cross-validation. In addition, model predictions for *G′*, thickness and specific curli fibre content were tested on an independent experimental test set comprising previously unseen PAS repeats with n = 4, 8 and 15 at 15 µM IPTG and unseen IPTG level, 10 µM, for the CsgA-PAS5 variant (12 measurements, *n* = 3 living hydrogels for each of the 4 PAS-IPTG conditions). For permeability, training was performed with data for n = 0, 1, 5, 10 and 20 grown at 15 µM IPTG (15 measurements, *n* = 3 living hydrogels for each of the 5 PAS variants), and independent testing was performed on unseen PAS repeats with n = 4, 8 and 15 at 15 µM IPTG (9 measurements, *n* = 3 living hydrogels for each of the 3 PAS variants).

#### Leave-One-Condition-Out (LOCO)

An experimental condition was defined as a unique parameter pair (*N*_PAS_, *C*_IPTG_). With 5 PAS levels and 3 IPTG levels, the dataset contains 15 distinct condition groups. In each fold of LOCO cross-validation, all replicates corresponding to one unique condition were omitted from the training set and held out exclusively as the test set (*k* = 15 folds). This quantifies the capacity of the model to predict living hydrogel properties at unmeasured design coordinates.

#### Leave-One-IPTG-Out (LOIO)

To evaluate model performance under complete chemical domain shifts^37^, all samples corresponding to a single IPTG concentration (e.g., all constructs induced at *C*_IPTG_ = 15 *μ*M across all PAS length variants) were removed from training and utilized as the test set (*k* = 3 folds). LOIO tests the model’s ability to interpolate or extrapolate along the chemical induction axis, demonstrating its robustness against out-of-distribution observations.

#### Manhattan distance

To quantitatively evaluate the inverse design framework, we measured parameter recovery accuracy by computing the Manhattan distance (ℓ₁-norm) in parameter space between the retrieved input coordinates (*PASn_pred_*, *IPTG_pred_*) and the ground-truth test coordinates (*PASn_true_*, *IPTG_true_*):

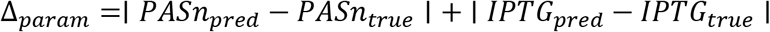

#### Statistical analysis

One-way ANOVA was used to compare multiple groups, and a two-way ANOVA for analyses involving multiple groups and two experimental factors. The specific statistical test, the number of replicates, the total number of observations as well as additional details are indicated in the corresponding figure and table legends. Unless stated otherwise, experiments were performed with *n* = 3 living hydrogels prepared from the same culture batch and grown and characterized independently. Statistical differences are reported as significant for *P* < 0.05 and non-significant for *P* ≥ 0.05.

#### Software

Statistical analyses, data plotting, figure formatting and assembly were performed using GraphPad Prism 10.2 (GraphPad Software). Rheological data was collected using the RheoCompass Software v1.32. OCT image acquisition was performed using ThorImage OCT v5.6. Image processing and analysis were performed using ImageJ v1.54i^51^ including the BioVoxxel toolbox v2.6^52^. Visualization and processing of 3D protein structures were performed using Jmol v16.1.1^49^. BioRender was used to prepare schematic representations. When required figures were assembled in Microsoft PowerPoint.

Regarding the predictive model implementation and evaluation, data preprocessing, array manipulation, and statistical analyses were performed in Python (v3.10)^57^ using NumPy (v1.24)^58^ and pandas (v2.0)^59^. Linear regression and GPR models were implemented using scikit-learn (v1.3)^60^. Tabular foundation modelling was conducted using TabPFN v2 (v8.0.3)^21^. All model training, prediction, and evaluation procedures, including the LOCO and LOIO cross-validation schemes, were implemented in Python.

During the manuscript preparation, the authors used Microsoft Copilot to improve readability and language. All AI assisted text was reviewed and edited by the authors, who take full responsibility for the content of the publication.

#### Computational Resources and Runtime

All computations were performed on a local workstation equipped with an Intel Core i9-14900K CPU (24 cores, 32 logical threads), an NVIDIA RTX 4500 Ada Generation GPU (24 GB VRAM), and 128 GB of RAM. The complete data-processing and model-evaluation pipeline, including cross-validation was executed in approximately 78 seconds. Because TabPFN operates as a pre-trained foundation model, pre-training time is excluded from the reported runtime.

## Data Availability Statement

All relevant data supporting the findings of this study are available within the Article and its Supplementary Information. The source code, trained models and datasets will be available upon acceptance at the GitHub repository: https://github.com/pgoo-dev/data-driven-elm-design.git.

The web-based platform for model deployment will be hosted at https://streamlit.io/ and the corresponding source code will be available through the same GitHub repository upon acceptance. Additional data present in this study is available from the corresponding author upon reasonable request.

## Acknowledgements

The authors thank Jan Becker for his guidance on rheometer operation and rheological data processing, Oliver S. Thomas for assembling the PAS1, PAS4, and PAS5 constructs, and Michael Backenköhler and Andrea Volkamer for their guidance during the initial exploration of predictive models. This work was supported by the European Research Council (ERC, grant agreement ID 101053857 - STEADY) and by the European Innovation Council (EIC, grant agreement ID 101070817 - LoopOfFun).

## Author contributions

G.M.-G., P.G. and W.W. conceptualized and designed the research, analysed the data, and wrote the manuscript with input from all authors. G.M.-G. designed and constructed the genetic variants used in this study, established the experimental methodologies and performed the experiments. P.G. developed and implemented the predictive modelling and inverse-design approaches, designed the machine-learning and validation workflows, curated and analysed the modelling data, and evaluated model performance. G.M.-G integrated the experimental and modelling results, prepared the figures, and assembled the manuscript. G.M.-G. and S.R. performed the spin-down quantitative Congo Red binding assay. S.R. and P.G implemented the Streamlit web-based platform. R.B. contributed to the development and validation of the mechanical characterization methodology and to the interpretation of rheological data. V.Z. contributed to the predictive models through methodological guidance and interpretation of modelling results. W.W. supervised the project and acquired funding. G.M.-G. and P.G. prepared the original draft of the manuscript. All authors read and commented on the manuscript.

## Supporting Information

Additional information can be found online in the Supporting Information section.

## Table of Contents (ToC) text

Rational design of engineered living materials (ELMs) is limited by the lack of quantitative relationships linking design parameters to material properties. In this study, a tabular foundation model informed by a small library of curli-based living hydrogels accurately predicts macroscopic material properties from genetic and process parameters. Property-guided design further enables identification of parameters for achieving ELMs with desired properties

## Data-driven predictive design of engineered living hydrogels

ToC figure

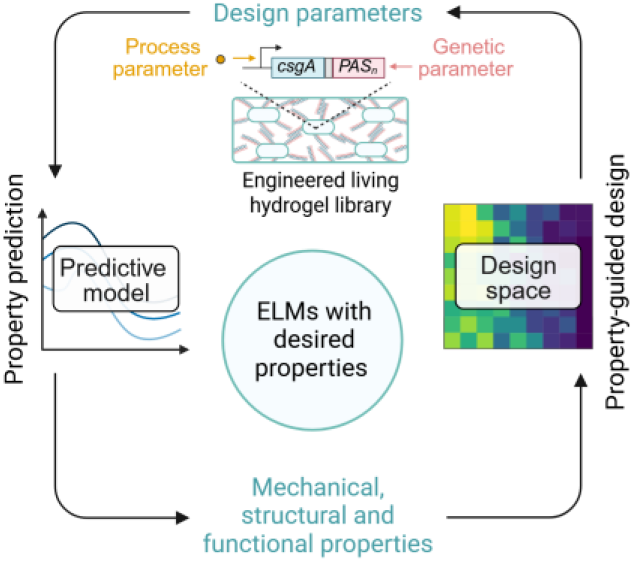

## References

1. Tang, T.-C. et al. Materials design by synthetic biology. Nat Rev Mater 6, 332–350 (2021).

2. Rodrigo-Navarro, A., Sankaran, S., Dalby, M. J., del Campo, A. & Salmeron-Sanchez, M. Engineered living biomaterials. Nat Rev Mater 6, 1175–1190 (2021).

3. Liu, A. P. et al. The living interface between synthetic biology and biomaterial design. Nat. Mater. 21, 390–397 (2022).

4. Gilbert, C. et al. Living materials with programmable functionalities grown from engineered microbial co-cultures. Nat. Mater. 20, 691–700 (2021).

5. McBee, R. M. et al. Engineering living and regenerative fungal–bacterial biocomposite structures. Nat. Mater. 21, 471–478 (2022).

6. Ge, C. et al. Engineered living glues secrete therapeutic proteins for treatment of inflammatory bowel disease. Nat Biotechnol 1–11 (2026) doi:10.1038/s41587-025-02970-9.

7. Manjula-Basavanna, A., Duraj-Thatte, A. M. & Joshi, N. S. Mechanically Tunable, Compostable, Healable and Scalable Engineered Living Materials. Nat Commun 15, 9179 (2024).

8. Duraj-Thatte, A. M. et al. Water-processable, biodegradable and coatable aquaplastic from engineered biofilms. Nat Chem Biol 17, 732–738 (2021).

9. Molinari, S. et al. A de novo matrix for macroscopic living materials from bacteria. Nat Commun 13, 5544 (2022).

10. Chen, A. Y. et al. Synthesis and patterning of tunable multiscale materials with engineered cells. Nature Mater 13, 515–523 (2014).

11. Nguyen, P. Q., Botyanszki, Z., Tay, P. K. R. & Joshi, N. S. Programmable biofilm-based materials from engineered curli nanofibres. Nat Commun 5, 4945 (2014).

12. Duraj-Thatte, A. M. et al. Programmable microbial ink for 3D printing of living materials produced from genetically engineered protein nanofibers. Nat Commun 12, 6600 (2021).

13. Praveschotinunt, P. et al. Engineered E. coli Nissle 1917 for the delivery of matrix-tethered therapeutic domains to the gut. Nat Commun 10, 5580 (2019).

14. Huang, J. et al. Programmable and printable Bacillus subtilis biofilms as engineered living materials. Nat Chem Biol 15, 34–41 (2019).

15. Jimenez, E. M. et al. Genetically Modifying the Protein Matrix of Macroscopic Living Materials to Control Their Structure and Rheological Properties. ACS Synth. Biol. 13, 3936–3947 (2024).

16. Liu, H., Chittur, P. K., Kornfield, J. A. & Tirrell, D. A. Cohesive Living Bacterial Films with Tunable Mechanical Properties from Cell Surface Protein Display. ACS Synth. Biol. 13, 3686–3697 (2024).

17. Duraj-Thatte, A. M. et al. Genetically Programmable Self-Regenerating Bacterial Hydrogels. Advanced Materials 31, 1901826 (2019).

18. Lopatkin, A. J. & Collins, J. J. Predictive biology: modelling, understanding and harnessing microbial complexity. Nat Rev Microbiol 18, 507–520 (2020).

19. Ramprasad, R., Batra, R., Pilania, G., Mannodi-Kanakkithodi, A. & Kim, C. Machine learning in materials informatics: recent applications and prospects. npj Comput Mater 3, 54 (2017).

20. Deringer, V. L. et al. Gaussian Process Regression for Materials and Molecules. Chem. Rev. 121, 10073–10141 (2021).

21. Hollmann, N. et al. Accurate predictions on small data with a tabular foundation model. Nature 637, 319–326 (2025).

22. Schmachtenberg, R. et al. Accelerated engineering and genetic programming of wood-based living composite materials. Materials Today 98, 103424 (2026).

23. Noda, R., Ichikawa, D. & Shibagaki, Y. Machine learning-based diagnostic prediction of minimal change disease: model development study. Sci Rep 14, 23460 (2024).

24. Yu, H. et al. Evaluating the effect of heart and respiratory rate measurement errors on the ability to predict the outcome of high flow nasal cannula therapy: a multi-centre study. Crit Care 29, 535 (2025).

25. Yan, Z., Yin, M., Chen, J. & Li, X. Assembly and substrate recognition of curli biogenesis system. Nat Commun 11, 241 (2020).

26. Schlapschy, M. et al. PASylation: a biological alternative to PEGylation for extending the plasma half-life of pharmaceutically active proteins. Protein Eng Des Sel 26, 489–501 (2013).

27. Breibeck, J. & Skerra, A. The polypeptide biophysics of proline/alanine-rich sequences (PAS): Recombinant biopolymers with PEG-like properties. Biopolymers 109, e23069 (2018).

28. Hidalgo, G., Chen, X., Hay, A. G. & Lion, L. W. Curli Produced by Escherichia coli PHL628 Provide Protection from Hg(II). Applied and Environmental Microbiology 76, 6939–6941 (2010).

29. Vidal, O. et al. Isolation of an Escherichia coli K-12 Mutant Strain Able To Form Biofilms on Inert Surfaces: Involvement of a New ompR Allele That Increases Curli Expression. J Bacteriol 180, 2442–2449 (1998).

30. Jumper, J. et al. Highly accurate protein structure prediction with AlphaFold. Nature 596, 583–589 (2021).

31. Mirdita, M. et al. ColabFold: making protein folding accessible to all. Nat Methods 19, 679–682 (2022).

32. Nikoumanesh, E., Jouaneh, C. J. M. & Poling-Skutvik, R. Elucidating the role of physicochemical interactions on gel rheology. Soft Matter 20, 7094–7102 (2024).

33. Olsen, B. D., Kornfield, J. A. & Tirrell, D. A. Yielding Behavior in Injectable Hydrogels from Telechelic Proteins. Macromolecules 43, 9094–9099 (2010).

34. Box, G. E. P. & Cox, D. R. An Analysis of Transformations. Royal Statistical Society. Journal. Series B: Methodological 26, 211–243 (1964).

35. Osborne, J. Notes on the use of data transformations. *Practical Assessment*, Research, and Evaluation 8, (2002).

36. Tukey, J. W. Exploratory Data Analysis. (Addison-Wesley Publishing Company, 1977).

37. Goodarzi, P., Schütze, A. & Schneider, T. Domain shifts in industrial condition monitoring: a comparative analysis of automated machine learning models. Journal of Sensors and Sensor Systems 14, 119–132 (2025).

38. Williams, P. C. & Sobering, D. C. Comparison of Commercial near Infrared Transmittance and Reflectance Instruments for Analysis of Whole Grains and Seeds. Journal of Near Infrared Spectroscopy 1, 25–32 (1993).

39. Bellon-Maurel, V., Fernandez-Ahumada, E., Palagos, B., Roger, J.-M. & McBratney, A. Critical review of chemometric indicators commonly used for assessing the quality of the prediction of soil attributes by NIR spectroscopy. TrAC Trends in Analytical Chemistry 29, 1073–1081 (2010).

40. Nawar, S. & Mouazen, A. M. Predictive performance of mobile vis-near infrared spectroscopy for key soil properties at different geographical scales by using spiking and data mining techniques. CATENA 151, 118–129 (2017).

41. Draper, N. R. & Smith, H. Applied Regression Analysis. (Wiley, 1998).

42. Rasmussen, C. E. & Williams, C. K. I. Gaussian Processes for Machine Learning. (MIT Press, Cambridge, Mass., 2008).

43. Gao, Y. & Cho, H. J. Quantifying the trade-off between stiffness and permeability in hydrogels. Soft Matter 18, 7735–7740 (2022).

44. Zunger, A. Inverse design in search of materials with target functionalities. Nat Rev Chem 2, 0121 (2018).

45. Sanchez-Lengeling, B. & Aspuru-Guzik, A. Inverse molecular design using machine learning: Generative models for matter engineering. Science 361, 360–365 (2018).

46. Gongora, A. E. et al. A Bayesian experimental autonomous researcher for mechanical design. Science Advances 6, eaaz1708 (2020).

47. Streamlit • A faster way to build and share data apps. https://streamlit.io/ (2021).

48. Bu, F., Dee, D. R. & Liu, B. Structural insight into Escherichia coli CsgA amyloid fibril assembly. mBio 15, e00419–24 (2024).

49. Hanson, R. M. Jmol – a paradigm shift in crystallographic visualization. J Appl Cryst 43, 1250–1260 (2010).

50. Wang, C., Lau, C. Y., Ma, F. & Zheng, C. Genome-wide screen identifies curli amyloid fibril as a bacterial component promoting host neurodegeneration. Proceedings of the National Academy of Sciences 118, e2106504118 (2021).

51. Schindelin, J., et al. Fiji: an open-source platform for biological-image analysis. Nat Methods 9, 676–682 (2012).

52. Brocher, J. biovoxxel/BioVoxxel-Toolbox: BioVoxxel Toolbox v2.6.0. (2023) doi:10.5281/zenodo.10050002.

53. Ostmann, K. et al. Bacterial Engineered Living Materials modulate Mechanosignaling in Mammalian Cells. 2024.10.29.620857 Preprint at 10.1101/2024.10.29.620857 (2024).

54. Duraj-Thatte, A. M. et al. Modulating bacterial and gut mucosal interactions with engineered biofilm matrix proteins. Sci Rep 8, 3475 (2018).

55. Buzza, K. M. et al. Modulation of Biofilm Formation and Permeability in Streptococcus mutans during Exposure To Zinc Acetate. Microbiology Spectrum 11, e02527–22 (2023).

56. Hubatsch, I., Ragnarsson, E. G. E. & Artursson, P. Determination of drug permeability and prediction of drug absorption in Caco-2 monolayers. Nat Protoc 2, 2111–2119 (2007).

57. Van Rossum, G. & Drake, F. L. Python 3 Reference Manual. (CreateSpace, Scotts Valley, CA, 2009).

58. Harris, C. R. et al. Array programming with NumPy. Nature 585, 357–362 (2020).

59. McKinney, W. Data Structures for Statistical Computing in Python. SciPy 2010 (2010) doi:10.25080/Majora-92bf1922-00a.

60. Pedregosa, F. et al. Scikit-learn: Machine Learning in Python. J. Mach. Learn. Res. 12, 2825–2830 (2011).

