## Supplementary Information for "Data-driven predictive design of engineered living hydrogels"

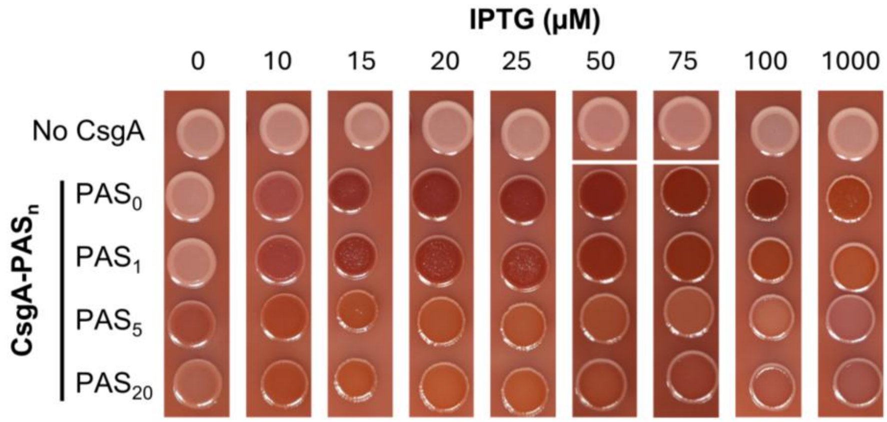

**Supplementary Figure 1. Optimization of IPTG concentrations**. CsgA-$PAS_n$ living hydrogels with n = 0, 1, 5 and 20 were used to scan a range of IPTG concentrations. For that, cultures of engineered bacteria were spotted on YESCA - Congo red agar plates containing IPTG concentrations ranging from 0 to 1000 µM. Curli fibre production was evaluated after 48 h growth at 25 °C.

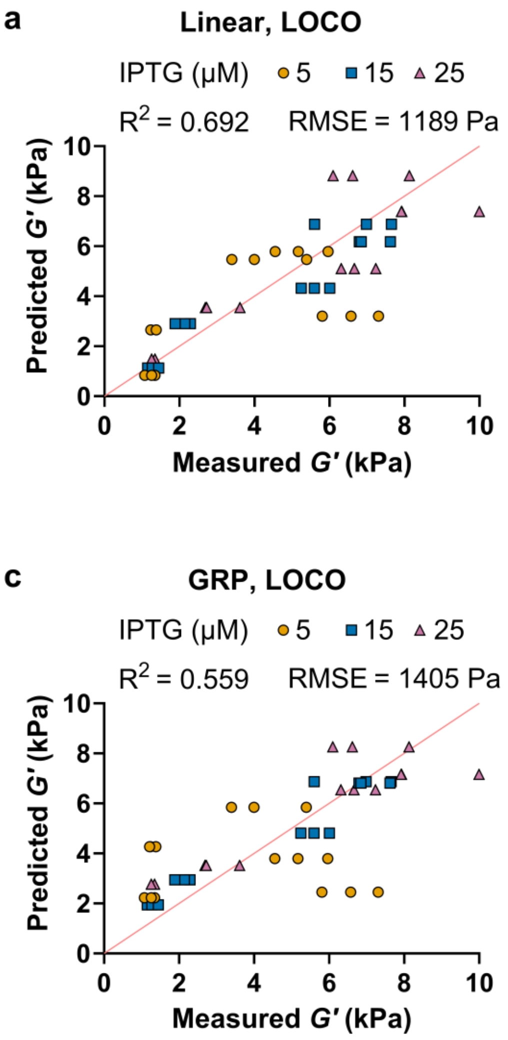

**Supplementary Figure 2. Predictive performance of linear regression and Gaussian process regression (GPR) for the storage modulus (*G′*) of living hydrogels under Leave-One-Condition-Out (LOCO) and Leave-One-IPTG-Out (LOIO) cross-validation**. The correlation between measured and predicted values was visualized using scatter plots for (**a, b**) linear regression and (**c, d**) GPR. Panels (**a, c**) correspond to LOCO cross-validation, in which individual PAS-IPTG combinations were withheld during training, and panels (**b, d**) correspond to LOIO cross-validation, in which an entire IPTG induction level was excluded from training and used for validation. The global $R^2$ and Root Mean Square Error (RMSE) for the different cross-validation strategies are displayed.

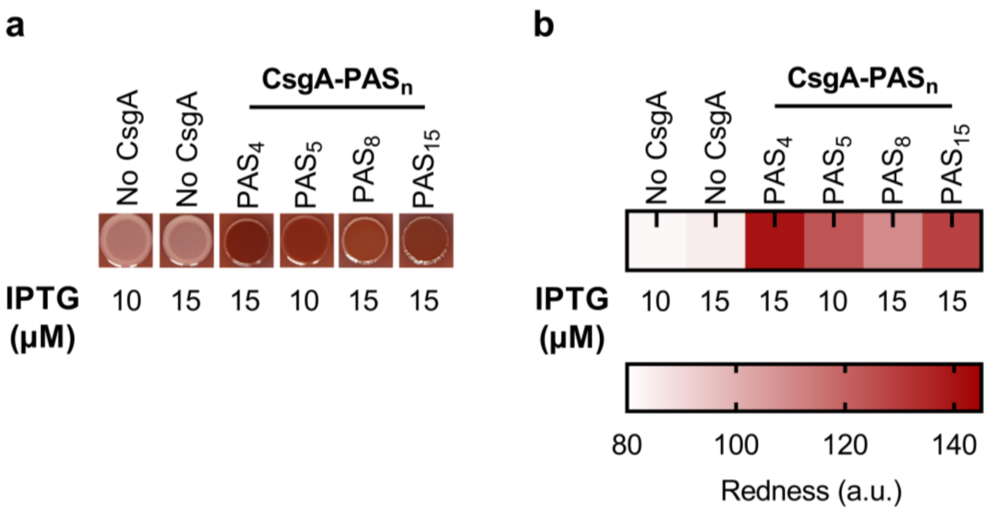

**Supplementary Figure 3. Engineered living hydrogel variants used as experimental validation dataset**. (**a**) Living hydrogels grown for 48 h at 25 °C using 15 µM IPTG and CsgA-PAS$_n$ variants with n = 4, 8 and 15 or using 10 µM IPTG with n = 5. (**b**) Quantification of living hydrogel redness to estimate curli fibre production.

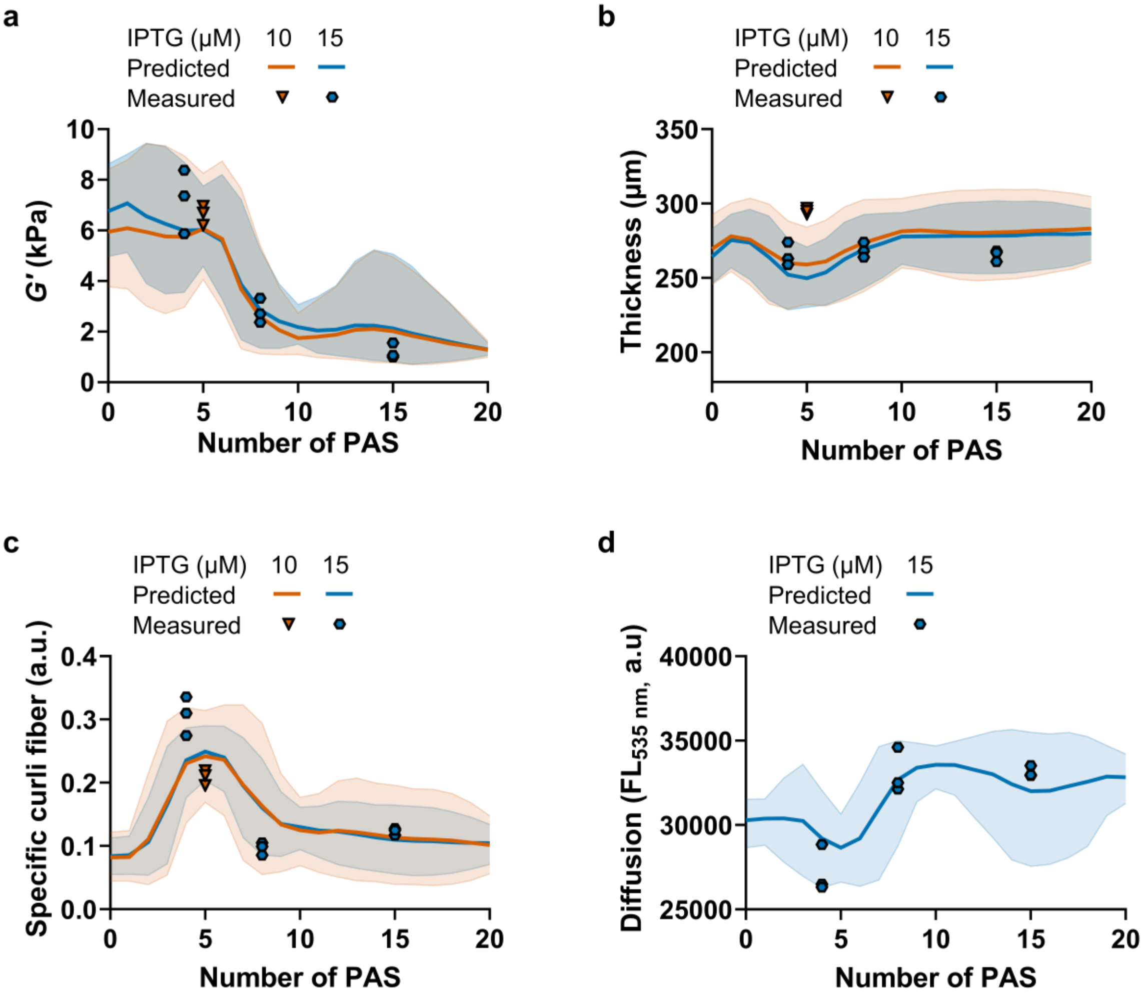

**Supplementary Figure 4. Visualization of independent test-set experimental values within the TabPFN predictive interval for the indicated living hydrogel properties**. The independent set used for experimental validation of predictions regarding (**a**) *G′*, (**b**) thickness and (**c**) specific curli fibre consisted of previously unseen PAS repeats (n = 4, 8 and 15) grown at 15 µM IPTG and unseen IPTG level (10 µM) for the CsgA-$PAS_5$ variant. (**d**) For permeability, the unseen PAS variants grown at 15 µM were used for experimental validation. Marker shapes represent experimental observations (for each PAS-IPTG condition *n* = 3 living hydrogels prepared from the same culture batch and grown and characterized independently). Solid lines represent the median of the predictive distribution. The shaded areas represent the uncertainty predictions, corresponding to the $5^{th}$ and the $95^{th}$ percentiles of the predictive distribution (90 % predictive interval).

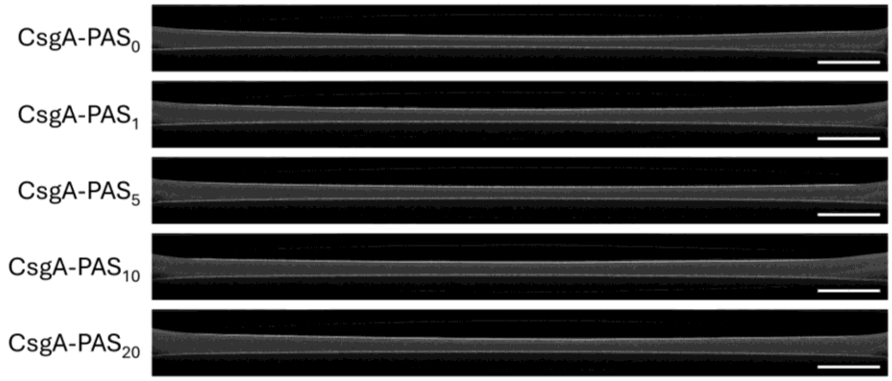

**Supplementary Figure 5. Representative cross-sectional images of living hydrogels obtained by optical coherence tomography (OCT)**. Living hydrogels were grown in plate inserts on YESCA agar plates containing 15 µM IPTG and incubated for 4 days at 25 °C. Then cross-sectional images were obtained by OCT, which were used to calculate the thickness of the living hydrogels using Fiji[1] including the BioVoxxel toolbox v2.6[2]. Scale bar: 1 mm.

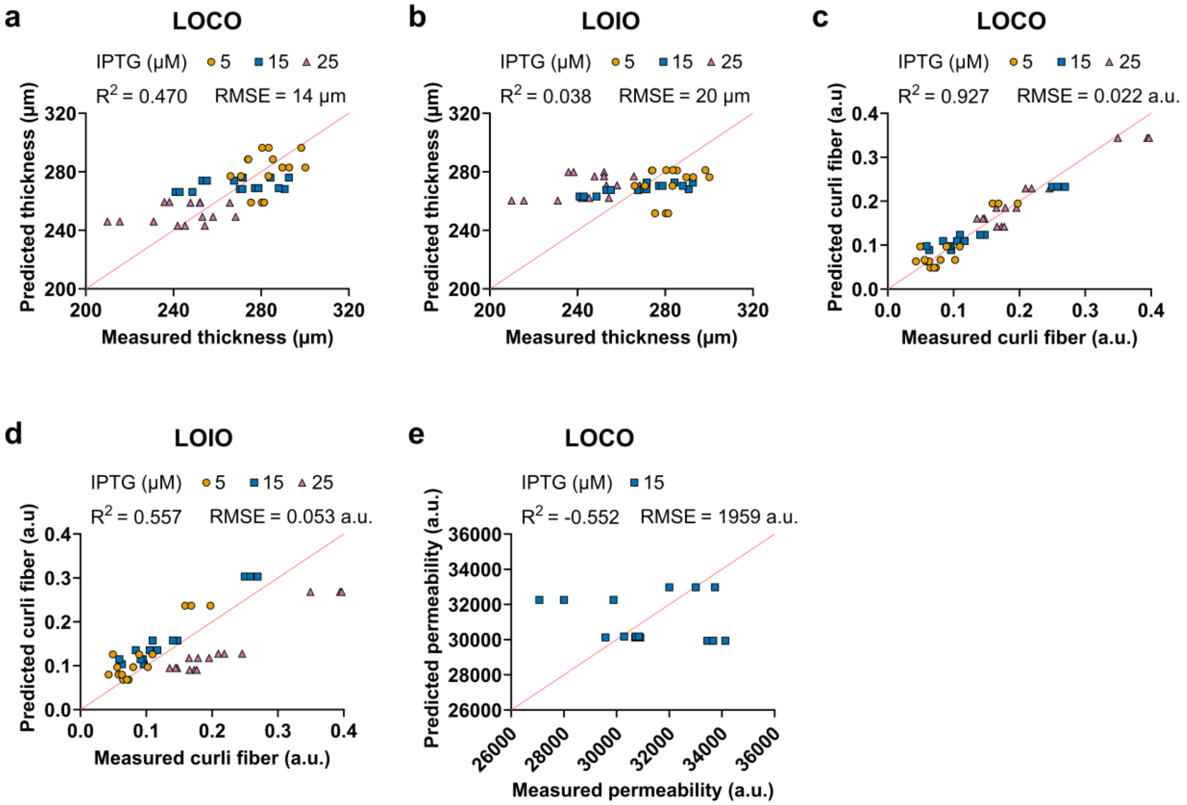

**Supplementary Figure 6. TabPFN predictive performance for the indicated living hydrogel properties under Leave-One-Condition-Out (LOCO) and Leave-One-IPTG-Out (LOIO) cross-validation**. The agreement between measured and predicted values is visualized using scatter plots for (**a, b**) thickness, (**c, d**) specific curli fibre and (**e**) permeability. Panels (**a, c, e**) correspond to LOCO cross-validation, in which individual PAS-IPTG combinations were withheld during training, whereas panels (**b, d**) correspond to LOIO cross-validation, in which an entire IPTG induction level was excluded from training and used for validation. The global $R^2$ and Root Mean Square Error (RMSE) for the different cross-validation strategies are displayed.

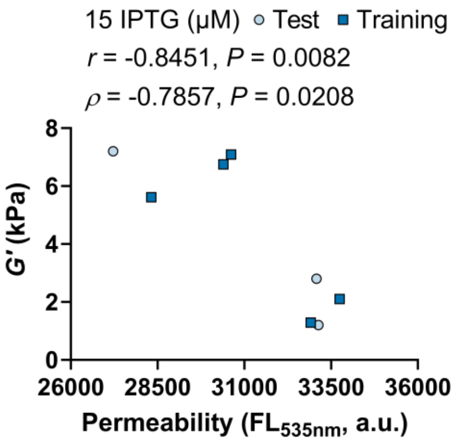

**Supplementary Figure 7. Relationship between living hydrogel permeability and storage modulus (*G'*)**. Each dot represents the mean of $n = 3$ living hydrogels measured for the CsgA-$PAS_n$ living hydrogels with n = 0, 1, 4, 5, 8, 10, 15 and 20 grown at 15 µM IPTG, where n = 4, 8 and 15 are part of the independent test set. Pearson's correlation coefficient (*r*) and Spearman's rank correlation coefficient (*ρ*) are reported in the panel.

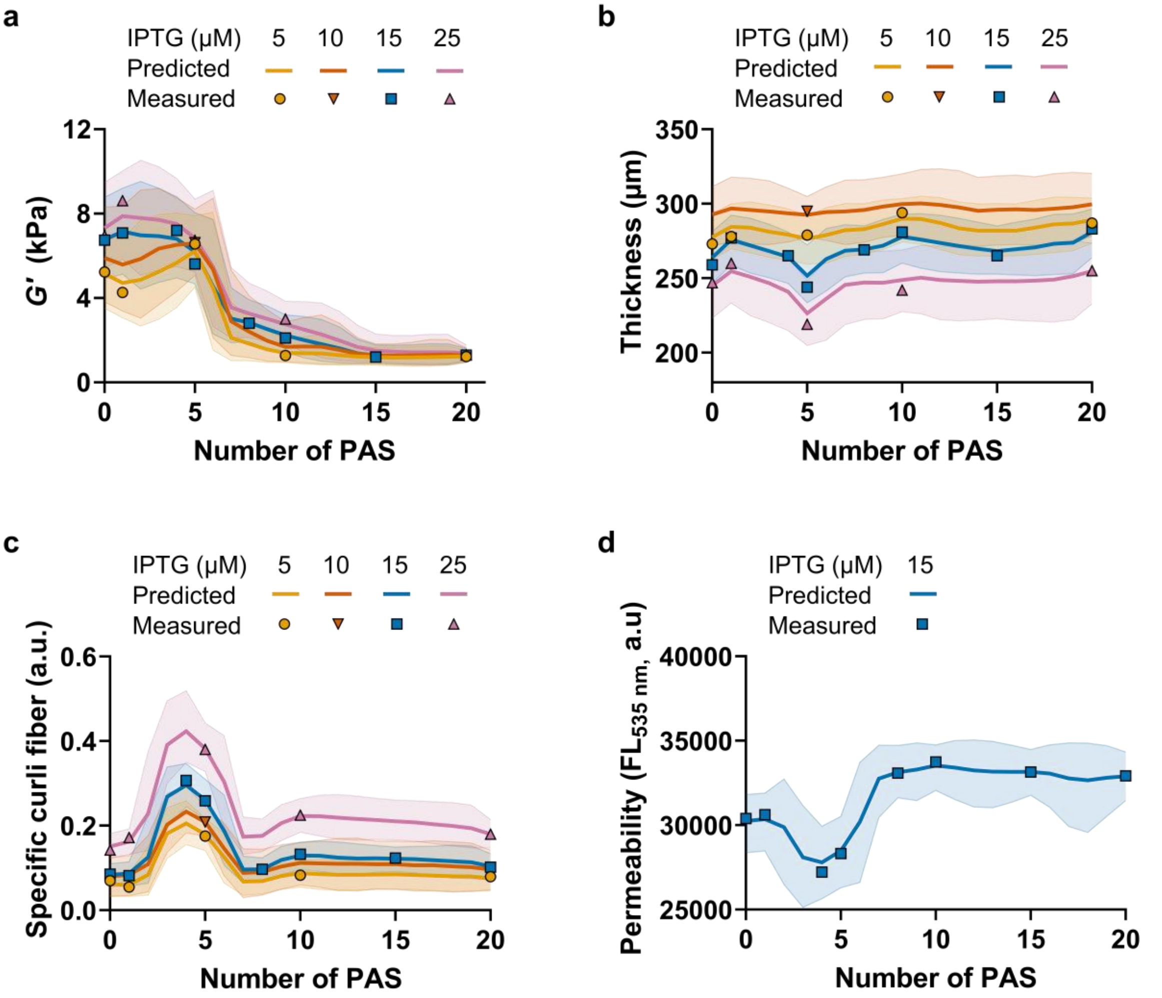

**Supplementary Figure 8. Prediction distribution of TabPFN with the complete dataset for the indicated living hydrogel properties**. TabPFN models were retrained with data hitherto used for experimental validation comprising CsgA-PAS$_n$ variants with n = 4, 8, 15 grown at 15 µM IPTG and n = 5 grown at 10 µM IPTG as well as the set previously used for training with n = 0, 1, 5, 10 and 20 grown at 5, 15, and 25 µM IPTG. The predictive interval was determined for (**a**) *G′*, (**b**) thickness, (**c**) specific curli fibre, and (**d**) permeability. Marker shapes represent the mean for *n* = 3 living hydrogels for each PAS-IPTG condition. Solid lines represent the median of the predictive distribution. The shaded areas represent the uncertainty predictions, corresponding to the to the 5$^{th}$ and the 95$^{th}$ percentiles of the predictive distribution (90 % predictive interval).

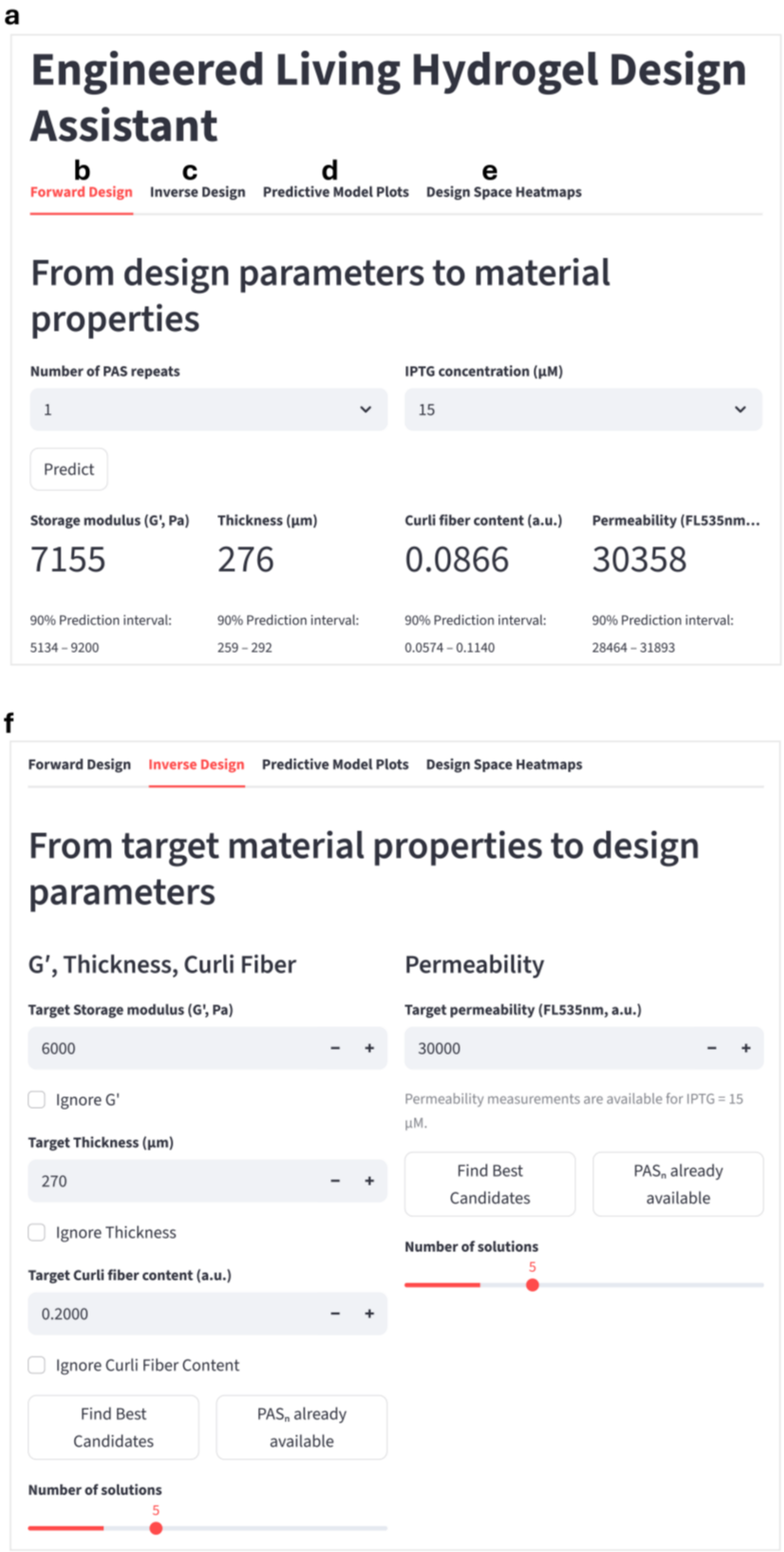

**Supplementary Figure 9. Web-based platform for model-assisted design of engineered living hydrogels**. (**a**) The interface integrates data-driven models for storage modulus ($G'$), thickness, specific curli fibre, and permeability. Users can perform (**b**) forward predictions to estimate material properties from selected design parameters or (**c, f**) inverse-design predictions to identify parameter combinations that achieve desired target properties. The platform additionally provides (**d**) visualizations of the predictions including their uncertainties (90% predictive interval), and (**e**) interactive heatmaps to explore the design space and identify parameter combinations that satisfy specified property requirements. This web-based platform will be hosted at https://streamlit.io/ and the corresponding source code will be available upon acceptance through the GitHub repository: https://github.com/pgoo-dev/data-driven-elm-design.git.

**Supplementary Table 1. DNA sequence of the engineered pCsgA-PAS$_n$ plasmids used in this study.** Where n denotates the number of repeats of the core PAS-encoding sequence fused to CsgA. pCsgA-PAS$_n$ with n= 0, 1, 4, 5, 8, 10, 15 and 20 were used in this study. Described plasmids were constructed in this study.

| Construct | DNA sequence |
|---|---|
| pCsgA-PAS$_n$:<br><br>n = 0, 1, 4, 5, 8, 10, 15, 20<br><br>backbone-P$_{trc}$-*lac* operon-RBS-*csgA*-*SapI*-*PAS$_n$*-*Nhe*I-Stop-backbone,<br><br>Cloning scars | TTCTGAAATGAGCTGTTGACAATTAATCATCCGGCTCGTATAATGTGTGGAATTGTGAGCGGATAACAATTTCAGAATTCAAAAGATCTTTTAAGAAGGAGATATACATATGAAACTTTTAAAAGTAGCAGCAATTGCAGCAATCGTATTCTCCGGTAGCGCTCTGGCAGGTGTTGTTCCTCAGTACGGCGGCGGTAACCACGGTGGTGGCGGTAATAATAGCGGCCCAAATTCTGAGCTGAACATTTACCAGTACGGTGGCGGTAACTCTGCACTTGCTCTGCAAACTGATGCCCGTAACTCTGACTTGACTATTACCCAGCATGGCGGCGGTAATGGTGCAGATGTTGGTCAGGGCTCAGATGACAGCTCAATCGATCTGACCCAACGTGGCTTCGGTAACAGCGCTACTCTTGATCAGTGGAACGGCAAAAATTCTGAAATGACGGTTAAACAGTTCGGTGGTGGCAACGGTGCTGCAGTTGACCAGACTGCATCTAACTCCTCCGTCAACGTGACTCAGGTTGGCTTTGGTAACAACGCGACCGCTCATCAGTACGGCTCTTCT(GCCTCTCCAGCTGCACCTGCTCCAGCAAGCCCTGCTGCACCAGCTCCGTCTGCTCCTGCT)$_n$GCCGCTAGCTGACTCGAGTAAGGATCT |

**Supplementary Table 2. Amino acid sequence of the engineered CsgA-PAS$_n$ fusion proteins**. Where n denotates the number of repeats of the PAS core sequence fused to CsgA. CsgA-PAS$_n$ with n= 0, 1, 4, 5, 8, 10, 15 and 20 were engineered and used in this study.

| **Construct** | **Amino acid sequence** |
|---|---|
| CsgA-PAS$_n$ :<br><br>n = 0, 1, 4, 5, 8, 10, 15, 20<br><br>CsgA-GSS-PAS$_n$-AAS | MKLLKVAAIAAIVFSGSALAGVVPQYGGGGNHGGGGNNSGP NSELNIYQYGGGNSALALQTDARNSDLTITQHGGGNGADVGQ GSDDSSIDLTQRGFGNSATLDQWNGKNSEMTVKQFGGGNGA AVDQTASNSSVNVTQVGFGNNATAHQYGSS(ASPAAPAPASPA APAPSAPA)$_n$AAS* |

**Supplementary Table 3. Ordinary two-way ANOVA test for the storage modulus (*G'*) of living hydrogels as a function of number of PAS repeats and IPTG concentration**. This test was performed with data for CsgA-PAS$_n$ variants with n = 0, 1, 5, 10 and 20 at 5, 15, and 25 µM IPTG (*n* = 3 living hydrogels for each PAS-IPTG condition, 45 measurements in total). Statistical significance was defined as $P < 0.05$.

| **Source of Variation** | **% of total variation** | ***P* value** | ***P* value summary** | **Significant?** | |
|---|---|---|---|---|---|
| Interaction | 7.252 | 0.0001 | *** | Yes | |
| Number of PAS | 81.75 | <0.0001 | **** | Yes | |
| IPTG (µM) | 6.523 | <0.0001 | **** | Yes | |
| **ANOVA table** | **SS** | **DF** | **MS** | **F (DFn, DFd)** | ***P* value** |
| Interaction | 21697845 | 8 | 2712231 | F (8, 30) = 6.081 | *P*=0.0001 |
| Number of PAS | 244610162 | 4 | 61152541 | F (4, 30) = 137.1 | *P*<0.0001 |
| IPTG (µM) | 19516120 | 2 | 9758060 | F (2, 30) = 21.88 | *P*<0.0001 |
| Residual | 13380139 | 30 | 446005 | | |

**Supplementary Table 4. Ordinary two-way ANOVA test for the specific curli fibre content of living hydrogels as a function of number of PAS repeats and IPTG concentration**. This test was performed with data for CsgA-PAS$_n$ variants with n = 0, 1, 5, 10 and 20 at 5, 15, and 25 µM IPTG (*n* = 3 living hydrogels for each PAS-IPTG conditions, 45 measurements in total). Statistical significance was defined as *P* < 0.05.

| **Source of Variation** | **% of total variation** | ***P* value** | ***P* value summary** | **Significant?** | |
|---|---|---|---|---|---|
| Interaction | 4.569 | 0.0002 | *** | Yes | |
| Number of PAS | 54.64 | <0.0001 | **** | Yes | |
| IPTG (µM) | 37.87 | <0.0001 | **** | Yes | |
| **ANOVA table** | **SS** | **DF** | **MS** | **F (DFn, DFd)** | ***P* value** |
| Interaction | 0.01539 | 8 | 0.001924 | F (8, 30) = 5.870 | *P*=0.0002 |
| Number of PAS | 0.1840 | 4 | 0.04601 | F (4, 30) = 140.4 | *P*<0.0001 |
| IPTG (µM) | 0.1276 | 2 | 0.06378 | F (2, 30) = 194.7 | *P*<0.0001 |
| Residual | 0.009830 | 30 | 0.0003277 | | |

**Supplementary Table 5. Ordinary one-way ANOVA test for the permeability of living hydrogels as a function of number of PAS repeats**. This test was performed with data for CsgA-$PAS_n$ variants with n = 0, 1, 5, 10 and 20 (*n* = 3 living hydrogels for each PAS condition, 15 measurements in total). Statistical significance was defined as $P < 0.05$.

| **ANOVA summary** | | | | | |
|---|---|---|---|---|---|
| F | 19.83 | | | | |
| *P* value | <0.0001 | | | | |
| *P* value summary | **** | | | | |
| Significant diff. among means (P < 0.05)? | Yes | | | | |
| R square | 0.8881 | | | | |
| **Brown-Forsythe test** | | | | | |
| F (DFn, DFd) | 0.8546 (4, 10) | | | | |
| P value | 0.5227 | | | | |
| P value summary | ns | | | | |
| Are SDs significantly different (P < 0.05)? | No | | | | |
| **ANOVA table** | **SS** | **DF** | **MS** | **F (DFn, DFd)** | ***P* value** |
| Treatment (between columns) | 56147159 | 4 | 14036790 | F (4, 10) = 19.83 | *P*<0.0001 |
| Residual (within columns) | 7078016 | 10 | 707802 | | |
| Total | 63225175 | 14 | | | |
| **Data summary** | | | | | |
| Number of treatments (columns) | 5 | | | | |
| Number of values (total) | 15 | | | | |

**Supplementary Table 6. Ordinary two-way ANOVA test for the thickness of living hydrogels as a function of number of PAS repeats and IPTG concentration**. This test was performed with data for CsgA-$PAS_n$ variants with n = 0, 1, 5, 10 and 20 at 5, 15, and 25 µM IPTG (*n* = 3 living hydrogels for each PAS-IPTG condition, 45 measurements in total). Statistical significance was defined as $P < 0.05$.

| **Source of Variation** | **% of total variation** | ***P* value** | **P value summary** | **Significant?** | |
|---|---|---|---|---|---|
| Interaction | 11.87 | 0.0011 | ** | Yes | |
| Number of PAS | 24.26 | <0.0001 | **** | Yes | |
| IPTG (µM) | 54.01 | <0.0001 | **** | Yes | |
| **ANOVA table** | **SS** | **DF** | **MS** | **F (DFn, DFd)** | ***P* value** |
| Interaction | 2407 | 8 | 300.9 | F (8, 30) = 4.519 | *P*=0.0011 |
| Number of PAS | 4919 | 4 | 1230 | F (4, 30) = 18.47 | *P*<0.0001 |
| IPTG (µM) | 10951 | 2 | 5475 | F (2, 30) = 82.24 | *P*<0.0001 |
| Residual | 1997 | 30 | 66.58 | | |

**Supplementary Table 7. Single-property inverse search retrieval performance**. For each held-out test condition, inverse search was performed targeting the mean $G'$. Reported candidates include the Top 3 retrieved input coordinate ($PAS_n$, IPTG). Parameter distance is calculated as the Manhattan ($\ell_1$) distance between the Top 1 retrieved input coordinates and the ground-truth test input parameters.

| **Test condition ($PAS_n$, IPTG)** | **Target $G'$, (Pa)** | **Top 3 candidates ($PAS_n$, IPTG) (1st; 2nd; 3rd)** | **Parameter distance** |
|---|---|---|---|
| (4, 15) | 7,202 | (3, 24); (3, 25); (3, 23) | 10 |
| (5, 10) | 6,624 | (0, 14); (0, 13); (0, 15) | 9 |
| (8, 15) | 2,799 | (14, 23); (14, 24); (14, 22) | 14 |
| (15, 15) | 1,203 | (20, 5); (20, 6); (19, 5) | 15 |

**Supplementary Table 8. Multi-property joint target search performance (3 properties).** For each held-out test condition, inverse retrieval was performed by jointly targeting the mean measured $G'$, thickness, and curli expression. Reported outputs include the target signature, Top 3 retrieved input coordinates ($PAS_n$, IPTG), predicted property values and Manhattan ($\ell_1$) parameter distance.

| **Condition ($PAS_n$, IPTG)** | **Target properties ($G'$, thickness, curli)** | **Top 3 candidates ($PAS_n$, IPTG) (1st; 2nd; 3rd)** | **Predicted $G'$ (Pa)** | **Predicted thickness (µm)** | **Predicted specific curli fiber content** | **Parameter distance** |
|---|---|---|---|---|---|---|
| (4, 15) | 7202 Pa, 265 µm, 0.3066 a.u. | (5, 10); (5, 11); (5, 9) | 6,071 | 257 | 0.2418 | 6 |
| (5, 10) | 6625 Pa, 295 µm, 0.2088 a.u. | (6, 5); (5, 5); (4, 5) | 5,631 | 275 | 0.1716 | 6 |
| (8, 15) | 2800 Pa, 269 µm, 0.0966 a.u. | (9, 15); (14, 18); (15, 19) | 2,413 | 273 | 0.1350 | 1 |
| (15, 15) | 1203 Pa, 265 µm, 0.1231 a.u. | (20, 22); (20, 21); (19, 21) | 1,373 | 268 | 0.1475 | 12 |

**Supplementary Table 9. Multi-property search with design-parameter constraints and preferences**. Multi-property inverse retrieval across four practical engineering scenarios integrating design parameter constraints (e.g., available CsgA-$PAS_n$ construct sets with n = 4, 8 and 15, or fixed IPTG concentrations) and preference weighting for lower IPTG levels or shorter PAS repeat lengths. Reported metrics include the reference condition, constraint descriptions, top 3 retrieved candidate ($PAS_n$, IPTG), predicted property values, and Manhattan ($\ell_1$) parameter distance.

| Scenario | Reference condition (PASn, IPTG) | Design parameter and property constraints | Top 3 retrieved candidates (1st ; 2nd ; 3rd ) | Predicted $G'$ (Pa) | Predicted thickness (µm) | Predicted curli fiber content | Parameter distance |
|---|---|---|---|---|---|---|---|
| High $G'$ + Lower IPTG (≤10 µM) and Short PAS (≤5) | (4, 15) | $G'$ ≥ 5000 Pa; **$PAS_n$** ∈ [0–5]; **IPTG** ∈ [5–10] µM; **lower IPTG-PAS** prioritized | (5, 6); (4, 7); (5, 7) | 6,130 | 267 | 0.2022 | 10 |
| Moderate $G'$ + experimental PAS Repeats {4, 8, 15} | (8, 15) | $G'$ ∈ [1500, 4000] Pa; **$PAS_n$** ∈ {4, 8, 15}; IPTG ≤ 15 µM; **lower IPTG** prioritized | (8, 6); (8, 7); (8, 8) | 2,217 | 281 | 0.1234 | 9 |
| Low $G'$ + moderate PAS length ([10–20]) and lower IPTG (≤15 µM) | (15, 15) | $G'$ ≤ 2000 Pa; **$PAS_n$** ∈ [10–20]; **IPTG** ∈ [5–15] µM; **lower IPTG** prioritized | (12, 13); (11, 14); (16, 14) | 1,977 | 280 | 0.1240 | 5 |
| Fixed IPTG = 10 µM + High/Moderate G' (Target Test Condition (5, 10)) | (5, 10) | **Fixed IPTG** = 10 µM; **$G'$** ≥ 4000 Pa; **Thickness** ≥ 250 µm | (3, 10); (6, 10); (4, 10) | 5,789 | 268 | 0.1708 | 2 |

**Supplementary Table 10. Predictive performance of TabPFN models for *G′*, thickness, specific curli fibre, and permeability with the complete dataset, including the dataset previously reserved for experimental validation**. TabPFN models were retrained with data hitherto used for experimental validation comprising CsgA-PASn variants with n = 4, 8, 15 grown at 15 µM IPTG and n = 5 grown at 10 µM IPTG as well as the set previously used for training with n = 0, 1, 5, 10 and 20 grown at 5, 15, and 25 µM IPTG. Model accuracy is reported for Leave-One-Condition-Out (LOCO) cross-validation, Leave-One-IPTG-Out (LOIO) cross-validation. Model accuracy is reported using Root Mean Square Error (RMSE), $R^2$ and the Ratio of Performance to Interquartile Distance (RPIQ). SD: standard deviation.

| **Property** | **LOCO cross-validation** | | | **LOIO cross-validation** | | |
|---|---|---|---|---|---|---|
| | **$R^2$** | **RMSE (mean ± SD)** | **RPIQ** | **$R^2$** | **RMSE (mean ± SD)** | **RPIQ** |
| *G′* | 0.759 | 1014 ± 763 Pa | 4.1 | 0.682 | 1264 ± 594 Pa | 3.5 |
| Thickness | 0.340 | 14 ± 8 µm | 1.7 | -0.608 | 28 ± 6 µm | 1.1 |
| Specific curli fibre | 0.834 | 0.028 ± 0.02 a.u. | 3.0 | 0.583 | 0.057 ± 0.019 a.u. | 1.9 |
| Permeability | 0.800 | 984 ± 451 a.u. | 3.1 | - | - | - |

### References

1. Schindelin, J. *et al.* Fiji: an open-source platform for biological-image analysis. *Nat Methods* **9**, 676–682 (2012).
2. Brocher, J. biovoxxel/BioVoxxel-Toolbox: BioVoxxel Toolbox v2.6.0. https://doi.org/10.5281/zenodo.10050002 (2023) doi:10.5281/zenodo.10050002.
